# Subtomogram averaging of helical filaments in cryo-electron tomography

**DOI:** 10.1101/2024.03.04.583254

**Authors:** Xiaojie Zhang, Julia Mahamid

**Author notes:** Lead contact: Julia Mahamid.

## Abstract

Helical filaments are essential macromolecular elements in cellular organization and dynamics. Recent advances in cryo-electron tomography allow faithful imaging of isolated or in-cell helical filaments. Here, we present a protocol to generate density maps at sub-nanometer resolution of helical filaments by subtomogram averaging, exemplified with isolated mumps virus nucleocapsids and their in-cell form as an extension of the method. We detail procedures from pre-processing of tilt series frames to refinement of reconstructed averages for streamlined data processing of helical filaments.

---

**For complete details on the use and execution of this protocol, please refer to Zhang et al.^1^**

## Before you begin

The protocol below describes the specific steps required for determining structures of mumps virus nucleocapsids as an example of heterogenous biological filaments with helical symmetry (raw micrographs in EMPIAR: 10751). The nucleocapsids were isolated from HeLa cells persistently infected with mumps virus after 1 hour of sodium arsenite stress treatment, imaged by cryo-electron tomography (cryo-ET) and computationally analyzed by subtomogram averaging (Figure 1). We have also used this protocol with slight modifications for determining the structure of nucleocapsids inside cells after 6 hours of potassium arsenate or sodium arsenite treatment.^1^

**Figure 1.**
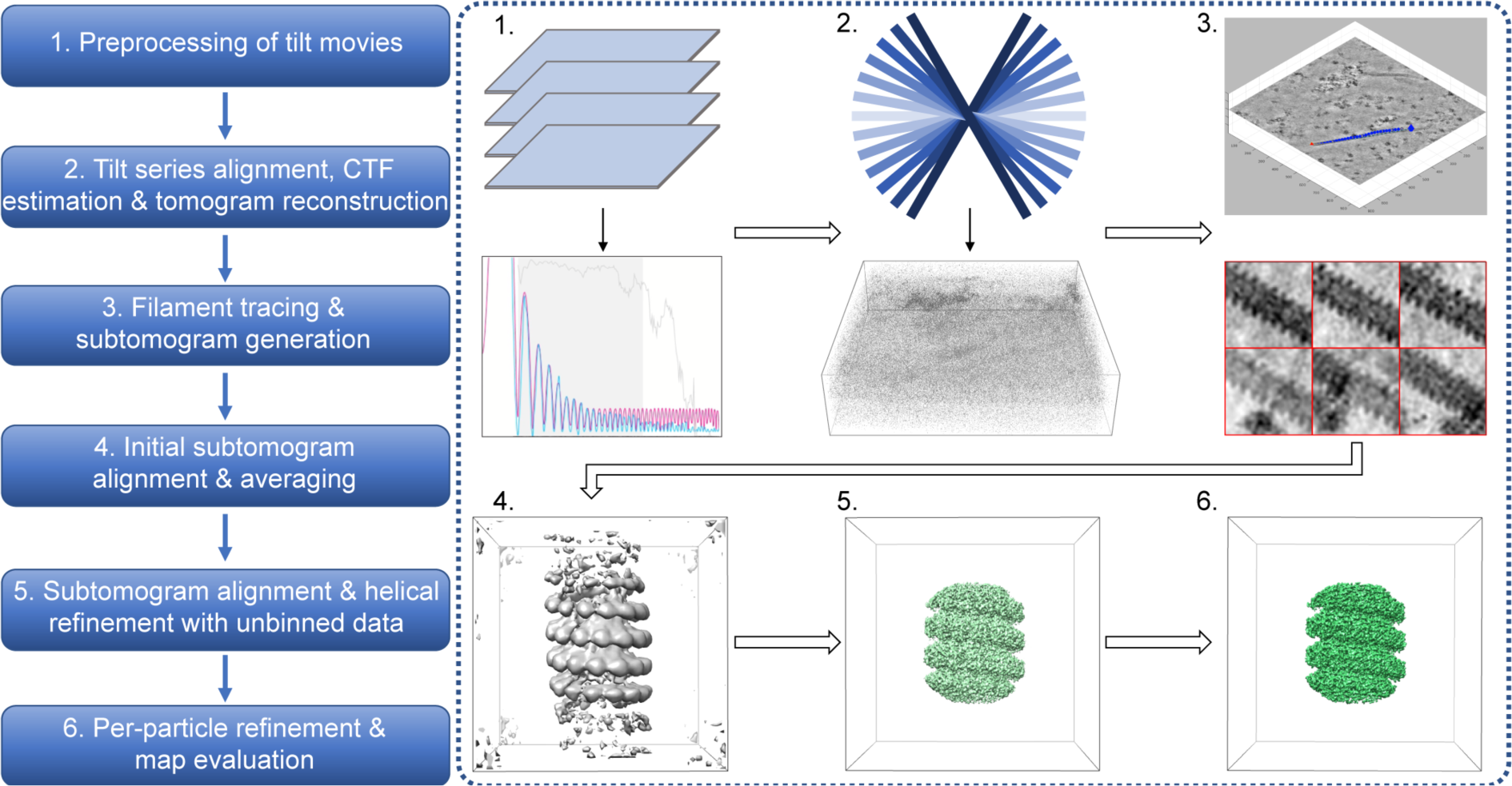
Overview of the pipeline for subtomogram averaging of filamentous helical assemblies described in this protocol.

### Preparation of cryogenically preserved helical assemblies

#### Timing: 6 – 8 hours

1. We isolated mumps viral nucleocapsids from HeLa cell lysate by pelleting at 13,000 *g* after clearing of cell debris at 500 *g*. The 13,000 *g* pellet enriched with nucleocapsids was subjected to heparin treatment in order to induce straightening of the nucleaocapsids.^1^ Straight nucleocapsids proved beneficial for obtaining high resolution maps.
2. Plunge freeze the isolated nucleocapsids.

a. Glow-discharge holey Quantifoil grids overlaid with an additional layer of continuous carbon (R2/1 + 2 nm C, Cu 200 mesh grid, Quantifoil Micro Tools). This additional layer of carbon may help to provide more uniformly oriented horizontal filaments on the grid surface, which helps in obtaining uniform ice thickness and in filament tracing.
b. Add Protein-A, 10-nm colloidal gold (Electron Microscopy Sciences) prior to freezing the grids. In our experiments, addition of 1.5 μl of 3 × concentrated fiducials to every 3 μl of nucleocapsid resuspension per grid still resulted in a low number of fiducials within the field of view of the tilt series. Therefore, tilt-series alignment was eventually performed with patch tracking (see sections below). Optimization of fiducial number might be beneficial for alignment of tilt-series with lower signal-to-noise acquired on thicker specimens.^2^

### Cryo-electron tomography data acquisition and computational setup

#### Timing: 10 – 20 hours

3. Acquire tilt series data on a microscope with the following or similar configuration: Titan Krios microscope operated at 300 kV (Thermo Fisher Scientific) equipped with a field-emission gun, a Quantum post-column energy filter (Gatan) and a K2 direct detector camera (Gatan). Record data using dose-fractionation mode in a dose-symmetric scheme^3^ with automated acquisition procedures as implemented in SerialEM software (v3.7.2)^4^, at a calibrated pixel size at the specimen level of 1.69 Å, with a defocus range of 2.5 to 3.5 µm, and a total dose of up to 120 e^-^/Å^2^ for 41 tilts. In total, we acquired 23 tilt series for the relatively straight isolated nucleocapsids. Out of these, 3 tilt series did not contain filaments, whereas the remaining 20 commonly contained a single filament. The filaments in the data must exhibit straight segments in order to achieve the final averages described in this protocol. Curved filaments are likely to exhibit larger local conformational heterogeneity, which in turn contributes to lower resolution reconstruction.^1^
4. Assess the data quality before going through the extensive data processing steps. For example, process a few tilt series in Warp and IMOD first. Overall, if the estimated maximum resolution of the inspected tilt series determined by the Contrast Transfer Function (CTF) fitting in Warp does not go beyond 5 Å and the reconstruction mean residual error is on average larger than 1 nm, we expect the highest achievable resolution of averages to be limited.
5. Ensure that the software and versions detailed in the Key resources table are installed on your workstation, server or high-performance-computing cluster. Ensure hardware with sufficient memory, CPU and GPU capacities. For example, we run Warp and M on a virtual Windows system including 2 Intel Xeon Gold 6226R CPU (total 32 cores, 2.9 GHz), 2 Nvidia Grid V100S GPUs (32 GB memory), and 128 GB RAM; most dynamo and MATLAB jobs were run on a server with Linux system including 2 AMD EPYC 7513 CPU (total 64 cores, 2.6 GHz), 2 Nvidia GeForce RTX 3090 GPUs (24 GB memory), and 256 GB RAM; RELION jobs were run on a cluster with multiple GPUs (e.g., 3090, A40).

## Key resources table

| REAGENT or RESOURCE | SOURCE | IDENTIFIER |
| --- | --- | --- |
| Deposited data |  |  |
| EM map for isolated nucleocapsids, majority class | Zhang et al.<sup>1</sup> | EMD-13133 |
| EM map for isolated nucleocapsids, minority class | Zhang et al.<sup>1</sup> | EMD-13136 |
| Model for isolated nucleocapsids, majority class | Zhang et al.<sup>1</sup> | PDB: 7OZR |
| Raw micrographs for isolated nucleocapsids | Zhang et al.<sup>1</sup> | EMPIAR-10751 |
| Software and algorithms |  |  |
| IMOD package | Kremer et al.<sup>5</sup> | https://bio3d.colorado.edu/imod/ |
| Warp 1.0.7 beta and 1.0.9 | Tegunov and Cramer<sup>6</sup> | http://www.warpem.com |
| M 1.0.9 | Tegunov et al.<sup>7</sup> | http://www.warpem.com |
| MATLAB R2019a and R2019b | MathWorks | https://www.mathworks.com |
| Dynamo 1.1.401 | Castano-Diez et al.<sup>8</sup> | https://wiki.dynamo.biozentrum.unibas.ch |
| TOM | Nickell et al.<sup>9</sup> | N/A |
| AV3 | Forster and Hegerl<sup>10</sup> | N/A |
| RELION 3.0 and 3.1 | Zivanov et al.<sup>11</sup> | https://github.com/3dem/relion |
| Chimera 1.13.1 | Pettersen et al.<sup>12</sup> | https://www.cgl.ucsf.edu/chimera/ |
| ChimeraX 1.1.1 | Pettersen et al.<sup>13</sup> | https://www.cgl.ucsf.edu/chimerax/ |
| dynamo2m | N/A | https://github.com/alisterburt/dynamo2m |

### Step-by-step method details

In the following, we provide a step-by-step guide for the generation of intermediate to high resolution electron microscopy (EM) density maps of helical filaments by subtomogram averaging (Figure 1). This includes preprocessing of tilt-series frame movies, tilt series alignment, CTF fitting, tomogram reconstruction, filament tracing, subtomogram generation, subtomogram alignment, averaging and classification at high to low data binning, per-particle refinement and finally EM map evaluation. To illustrate these steps, we use the example dataset of mumps virus nucleocapsids isolated from HeLa cells after 1 hour of sodium arsenite stress treatment.^1^ The data comprises 20 tilt series, each containing 1-2 filaments, acquired at a pixel size of 1.69 Å (EMPIAR-10751).

### Preprocessing of tilt movies

#### Timing: 1 – 2 hours

Pre-process the raw frame movies of individual tilts in 2D mode in Warp and generate tilt series stacks of the averaged images for tilt series alignment and tomogram reconstruction. We recommend referring to the original publications^6,7^ and the software website (http://www.warpem.com/warp/?page_id=378) for further details about the settings for Warp.

1. Preprocess the raw frame movies of individual tilts in 2D mode.

a. Provide all frame movies and the corresponding gain reference image from a single microscopy session in a single folder (e.g. named “frames”). This will constitute the folder for Warp processing, and all output files from Warp will be by default written into this “frames” folder. Provide all mdoc files (metadata text files for each tilt series written by SerialEM during the acquisition) in a separate folder at the same level as the “frames” folder. The names of the mdoc files will be taken as prefixes for tomograms in the next steps. Therefore, rename the mdoc files at this step if you wish to name the tomograms in a certain manner.
b. Launch Warp: execute the Warp.exe in the installation directory of Warp and add a desktop shortcut for future use.
c. Set up input parameters for processing (Figure 2).

i. Under the **Input** section, first locate the input “frames” folder. Then click *.tif to select the **TIF** option, set the pixel dimensions as 3708 x 3838 px (for K2 data), and select int16 to match the raw, unbinned frame images. Set the pixel X/Y as 1.6938 Å according to the unbinned pixel size of the data. Leave the **Bin** and **Dose** as default: 0.00x (unbinned) and 0.00 e/Å^2^/frame.
ii. Under the **Preprocessing** section, set the correct path to the gain reference image (.dm file in this case; this needs the “Flip Y axis” operation). For the example dataset, additionally select “Flip X axis” and “Transpose” because the raw images were saved as “r/f = 3” in relation to the gain reference image (read from the header of tif images; equivalent to 270° counterclockwise rotation of frame images relative to the gain reference image). Warp version 1.0.9 also allows input of a *.txt or *.tif defect map file to correct defects in camera sensors, which will be corrected along with the gain correction. **<u>Troubleshooting 1</u>**
iii. Under the **CTF** part of the **Preprocessing** section, set the power spectrum size in the **Window** value as 512 px (a relatively small window). To start with, set the spatial frequency **Range** to 25.0-5.0 Å (0.14-0.68 Ny, unbinned pixel size of 1.6938 Å) to test the CTF fitting. Adjust this range to include more of the lower or higher frequencies based on the quality of the CTF fitting test with a few movies before applying the range to all movies (switch to the **Fourier Space** tab and click **Process Only This Item’s CTF** to see whether the fitting range represented by the grey rectangle is optimal to the signal in your data). A range of 22.0-5.0 Å (0.15-0.68 Ny) worked well for the example data. Select the **Use Movie Sum** option as the fitting usually works better with the power spectrum of the movie average given the low dose per frame image in tilt-series data. Set the **Voltage** and **Cs** (spherical aberration) values as 300 kV and 2.70 mm respectively based on the microscope configuration. Set the **Amplitude** value to 0.07 (the fraction of amplitude contrast in the CTF; 0.07-0.10 recommended for cryo-electron microscopy (EM) data). Set the **Defocus** range as 1.0-4.0 µm based on the acquisition defocus range, with some margin.
iv. Under the **Motion** part of the **Preprocessing** section, set the **spatial frequency range** for the motion estimation to 67.8-13.6 Å (0.05-0.25 Ny) and the **B-factor** to -500 Å^2^.
v. Under the **Models** section, set the **Defocus** model resolution as 2x2x1, with the first two defining the spatial resolution and the third as temporal resolution for the defocus estimation (done on the sum of frames due to insufficient signals in a single frame). The recommended settings in the Warp manual 5x5x1 may also work depending on the unbinned pixel size of the data. Set the **Motion** resolution to 5x5x8 for this dataset, with the first two values defining the spatial resolution and the third defining the temporal resolution (8 frames per movie in the example dataset). As the manual suggests, set a temporal resolution value that matches the overall electron dose per pixel if a low dose rate (<1 e/px/frame) is used. The first two spatial resolution parameters can also be set to 1, if the signal in a single tilt movie average is suspected to be too low for alignment.
vi. Skip the **Pick Particles** option in this case.
vii. Leave the **Output** section as default. Do not skip any frames in this case. The aligned movie **Average** will be written out.
d. Start processing and check the results. With the parameters set in the previous steps, click the **START PROCESSING** button to start and the button will be switched to **STOP PROCESSING** once the procedure starts. Click to stop processing if parameters need be changed or the processing is completed. In this case, we only used one GPU but Warp will try to use all compatible GPUs available on the system. Therefore, deselect the unwanted GPUs from the upper right corner of the window bar. Warp provides an overview tab to show the processing status of all movies and plots for the defocus value, the estimated resolution, and the average motion per frame in the first 1/3 of the processed movies. On the left side of each plot, a histogram provides additional evaluation of the data quality.
e. When processing is done, one can check the processing results of individual tilt movies by clicking on the dots in each plot in the Overview tab and switching to the **Fourier & Real Space** tab, or directly in the Fourier & Real Space tab, clicking through the processing status bar (green/blue color in most cases) at the bottom of this tab (with the left and right arrow keys on the keyboard). For fine-tuning the CTF fitting parameters for individual tilt movies, one can use the **PROCESS ONLY THIS ITEM’S CTF button** after adjusting the spatial frequency range under the CTF section. If this is done, the previously processed items with different settings will change their processing status to **Outdated** in yellow. Changing the spatial frequency range back to the previous values after individual tilt movies are fine-tuned should change the process status of the majority movies back to green for proceeding to the next step. One can set the filter thresholds in all plots in the Overview tab so that movies with fitted parameters out of the selected range can be filtered out when proceeding with the Warp pipeline. Alternatively, one can manually exclude individual tilt movies by clicking on the check box in front of the name of each tilt movie at the navigation bar in the Fourier & Real Space window (“-” means automatic filters). In this example (Figure 2), movies were not excluded at this stage.
2. Export tilt series stacks for IMOD. In this pipeline, we use Warp to assemble tilt series stacks from tilt movies generated in the last step, align these stacks in IMOD, and returned to Warp to perform refinement of the CTF estimation for the tilt series and reconstruct them into tomograms. For assembling the tilt series, switch Warp to tilt series mode by choosing the **TomoSTAR** input format at the top of the window. In this mode, click on the **Import tilt series from IMOD** button at the Overview tab and a dialog pops up for choosing the path to the folder containing the mdoc files written by SerialEM and the path to the original input tilt movies (“frames” folder). Leave the pixel size as 1.6938 Å and dose per tilt as 0 e/Å^2^. Warp reads the dose value from the mdoc files. Uncheck the **Invert tilt angles** in the Warp version 1.0.7 beta (option changed to invert tilt angles by default in the Warp version 1.0.9). Click on **CREATE STACKS FOR IMOD**. Warp will write tilt series stacks into separate folders with the same name as the mdoc name for each.

**Figure 2.**
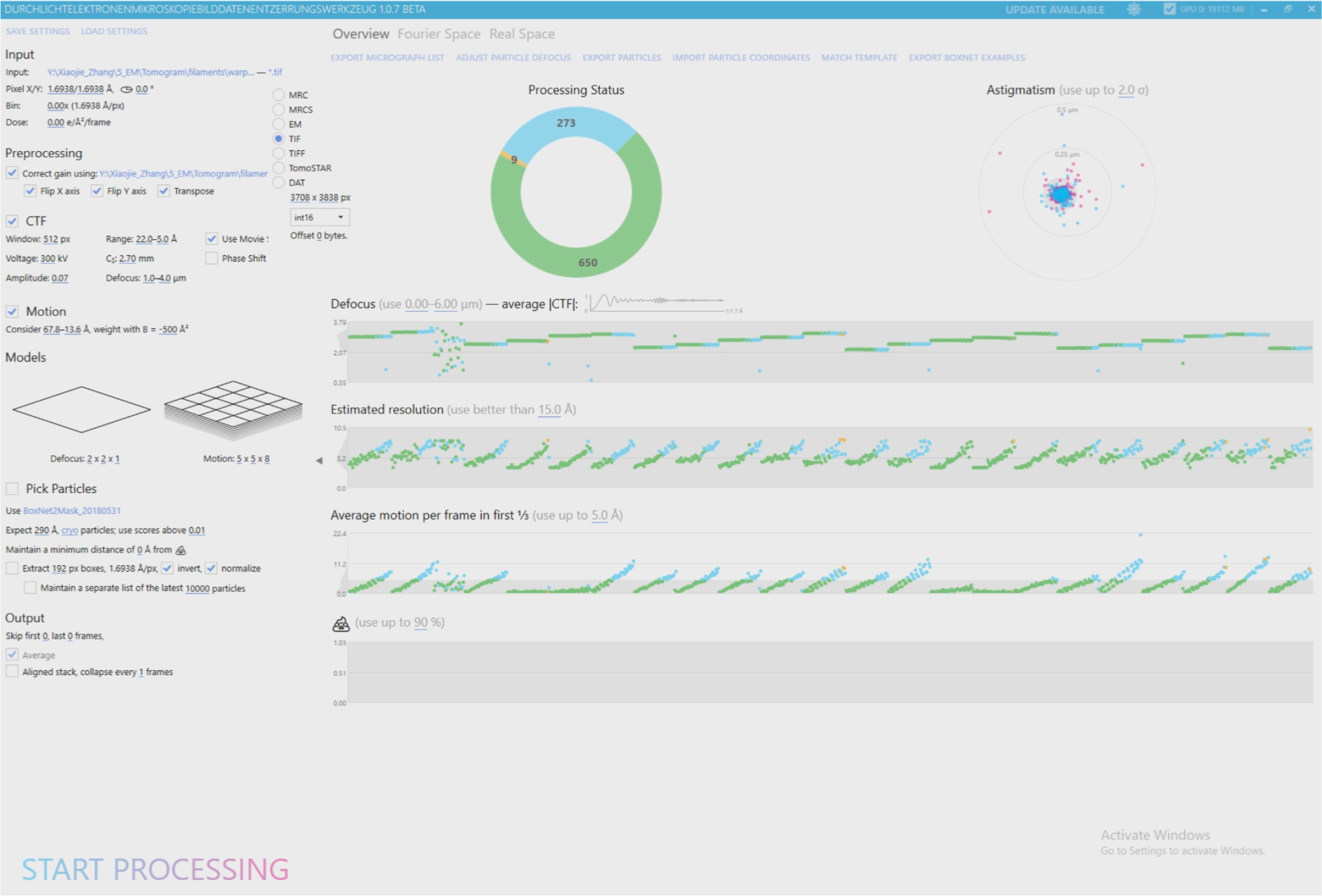
Screenshot of the Warp image pre-processing graphical user interface (GUI) and parameters used for processing the example data comprised of 20 tilt series (raw micrographs in EMPIAR-10751). Initially, 23 tilt series acquired during a single microscopy session were processed (shown in this screenshot), 3 of which were not processed further as they did not contain filaments.

### Tilt series alignment, CTF estimation and tomogram reconstruction

#### Timing: 7 hours

Align tilt series generated by Warp with Etomo in the IMOD package (version 4.9.4 described here). Processing of the tilt-series until the fine alignment step (included) is sufficient to allow Warp to import the alignment parameters from IMOD. Continue with CTF estimation for tilt series and reconstruct tomograms in Warp to the desired binning size for filament tracing in the next steps. For further information on processing in Etomo, we recommend referring to the software website (https://bio3d.colorado.edu/imod/doc/etomoTutorial.html).

3. Tilt series alignment with the Etomo module. Perform Etomo processing in the imod subfolders containing the individual tilt series written by Warp (Figure 3A). Leave parameters as default unless specified otherwise in the following steps.

a. With a Windows command prompt or Linux terminal, navigate to the subfolder for the first tilt series, and type “etomo” to launch the software. Select **Build Tomogram** to enter the setup interface for single tomogram reconstruction. Specify the tilt series to be reconstructed in **Dataset name**. Select cryoSample.adoc in **Templates** and set **Axis Type** as Single axis while keeping **Frame Type** as Single frame. Click on **Scan Header** to extract the **Pixel size** and **Image rotation** from the input data. Set **Fiducial diameter** as “0” as no fiducial markers will be used for tracking with this dataset. Select the “Extract tilt angles from data” option under the **Axis A** section. Click on the **View Raw Image Stack** button to inspect whether some views of the tilt series need be excluded at this point due to bad quality, e.g. views of areas largely covered by ice. Excluding such views at this Etomo reconstruction step instead of doing so within Warp avoids possible mismatch between Warp and Etomo reconstruction and the need of removing corresponding tilts by editing each mdoc file). Continue to **Create Com Scripts** to start processing.
b. At the **Pre-processing** window, click on the “Find X-rays (Trial Mode)” button to erase defects or “hot pixels” from tilt images. Click on “Create Fixed Stack” to create a fixed stack and continue to use it by clicking “Use Fixed Stack”. Click on “Done” to proceed.
c. At the **Coarse Alignment** window, first click on the “Calculate Cross-Correlation” button to align the tilt series. Then in the **Newstack** section, set the “Coarse aligned image stack binning” to 4 (6.78 Å/pix) and select the “Reduce size with antialiasing filter” and “Float intensities to mean”. Deselect “Convert to bytes”. Click on “Generate Coarse Aligned Stack” to generate a new binned stack for the next step. Click on “Done” to proceed.
d. At the **Fiducial Model Generation** window, select “Use patch tracking to make fiducial model” option and set the **Size of patches** to 150 by 150 pixels and **Iterations** as 4 (the maximum value). Click on “Track Patches” to generate the patch fiducial model. Click on “Done” to proceed.
e. At the **Fine Alignment** window, leave settings at the **General** tab as default and go to the **Global Variables** tab and set the “Rotation Solution Type” to “One rotation”, “Magnification Solution Type” to “Fixed magnification at 1.0”, “Tilt Angle Solution Type” to “Fixed tilt angles” (tilt angles pre-calibrated) and “Distortion Solution Type” to “Disabled”, respectively. Click on “Compute Alignment” and assess the alignment quality by reading the values corresponding to “Ratio of total measured values to all unknowns” and “Residual error mean and sd” in the Project Log. Aim for a residual error mean value smaller than 1 pixel if possible. For example, the 3^rd^ tilt series in this dataset resulted in an initial value of 1.150 pixels (sd: 1.018 pixels). The residual score was reduced to 0.756 pixels (sd: 0.532 pixels) after removal of a few bad contours (use “View/Edit Fiducial Model”). Right click at the Fine Alignment window to further open the log file with the “Align log file” option. Check the mean residual of each tilt view under the **Solution** tab in the log file in order to decide whether some views with large residual errors should be excluded for improving alignment, or use the “robust fitting” option, both of which are possible under the General tab. Click on “Done” to proceed.

**Figure 3.**
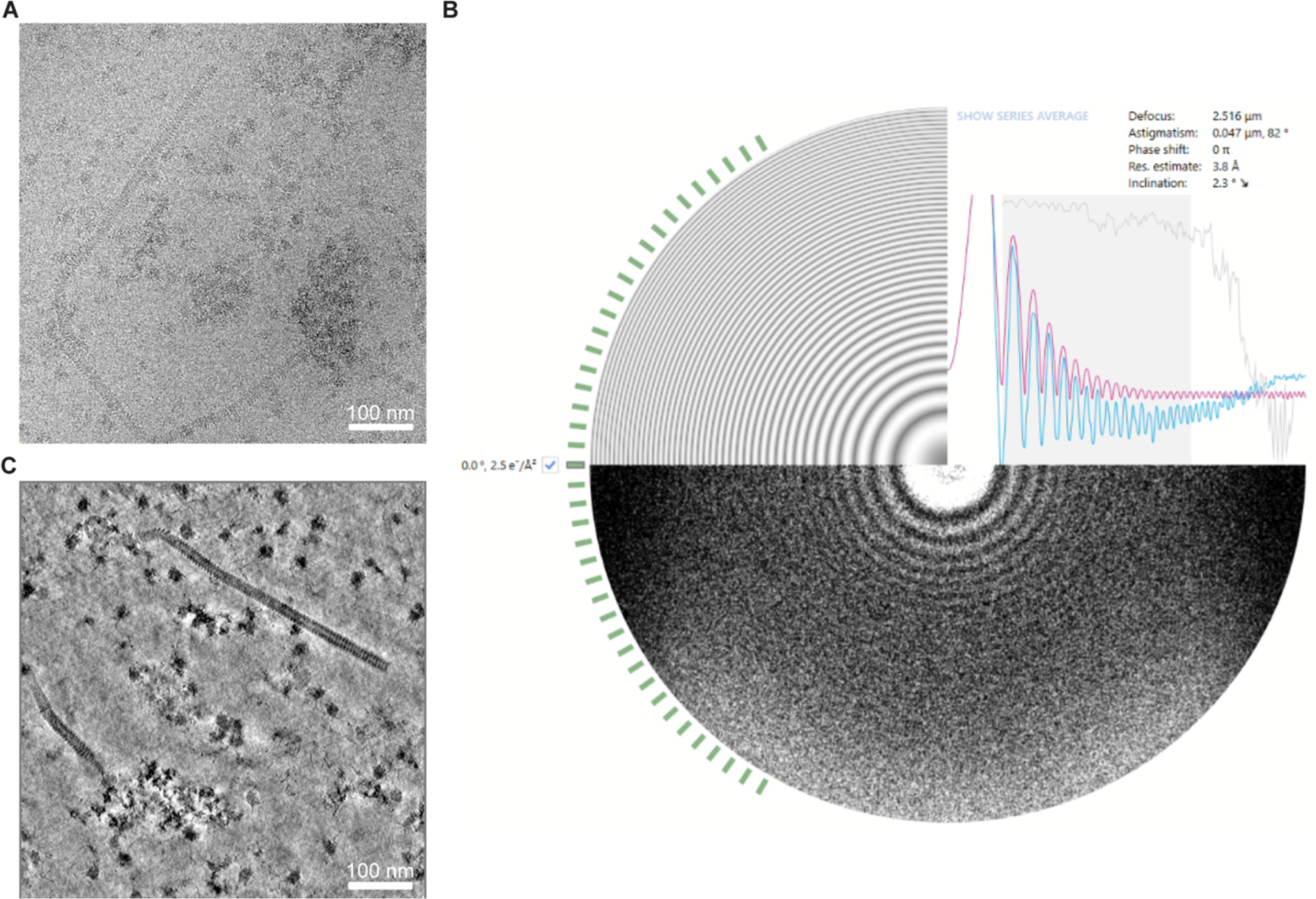
Tilt series CTF estimation and tomogram reconstruction in Warp. (A) A tilt image from one of the tilt series in the example data. (B) A screenshot from the Warp GUI showing the refined CTF processing result of the full tilt series in (A; the screenshot shows details for the 0-degree tilt) after importing alignment parameters determined in IMOD. The CTF fitting quality is normally expected to be worse at higher tilt angles compared to the ones at lower angles. (C) A tomographic slice of the tilt series shown in (A) following Warp reconstruction and deconvolution. The tomogram was deconvolved with the default parameters during the Warp reconstruction procedure for better visual inspection and manual filament tracing in the next step. The filament is curved, but its relatively straight segments can be used for subtomogram averaging. *For Warp reconstruction, the following steps f-j are not necessary. Warp makes use of the alignment parameters generated in Etomo up to this point. Proceed with the next steps for the first tilt series until the final tomogram is generated as a quality and consistency check*.
f. Skip the **Tomogram Positioning** step. While this step can help in reducing the reconstructed file size and providing easier visualization for tilted specimens, the transformation applied in Etomo can create a mismatch of particle positions as downstream processing is performed on untransformed reconstruction. Click “Done” to proceed.
g. At the **Final Aligned Stack** window, only use the Create tab. Enable “Use linear interpolation”, align image stack with a binning factor of 4, and enable “Reduce size with antialiasing filter”. Click on “Create Full Aligned Stack”. Click on “Done” to proceed.
h. At the **Tomogram Generation** window, use the “Back Projection” algorithm. Set the Tomogram thickness in Z to 1000 (in unbinned pixels). Radial Filtering is used here by default. Click on “Done” to proceed.
i. At the **Post-processing** window, deselect the “Convert to bytes” option under the **Scaling** section and select the “Rotate around X axis” option under the **Reorientation** section. This should ensure correct handedness in the reconstruction given the microscope setting in the example dataset. Click on “Done” to proceed.
j. Skip the **Clean Up** step for now and click on “Done” to finish reconstruction of the tilt series. Intermediate files of large sizes can be cleaned altogether when necessary.

**Optional:** Reconstructing tomograms in IMOD batch mode saves time especially with a larger dataset (https://bio3d.colorado.edu/imod/doc/batchGuide.html). Complete the reconstruction of one tomogram as detailed in this section and save the parameters in a “batch directive file” by using the **Export** option in the Etomo interface. In the batch reconstruction interface, set it as the “Starting directive file” and select “cryoSample.adoc” in the System template box. Import all the remaining tilt series in the “Stacks” tab, check parameters in the “Dataset Values” tab and set it to “Stop after Fine alignment” and press “Run” to start. One can assess the alignment results and adjust for individual tilt series if needed. Alternatively, one can use AreTomo^14^ for automatic batch alignment and reconstruction of tomograms with GPU environment available. Warp also accepts alignment parameters/files generated by AreTomo.

**Note**: We recommend to complete reconstruction of one tomogram per batch of data, for determining optimal parameters for tomogram reconstruction (e.g. Z dimension size), assessing the data quality, as well as checking the consistency between IMOD and Warp results.

4. Refinement of CTF estimation for tilt series and tomogram reconstruction in Warp.

a. Re-open Warp and click the **IMPORT TILT SERIES FROM IMOD** button to import the alignment results from IMOD in the tomostar mode. Specify the folders for mdocs, original frame movies, and IMOD processing results (select the subfolder named “imod” created by Warp). Due to a difference in handling tilt angle direction in Warp compared to IMOD, there is one option to invert tilt angle when importing tilt series from IMOD. When the example data were processed with the Warp version 1.0.7 beta, the tilt angle convention was set inversely to that in IMOD. Thus, additional flipping of subtomogram averages was necessary to produce the correct map handedness. If all IMOD processing results are found, check boxes in the “Aligned” column will automatically be ticked for all tilt series. Warp requires the xf and taSolution.log files in each imod reconstruction folders. Click on **IMPORT** to continue. **<u>Troubleshooting 2</u>**
b. Specify the **Unbinned tomogram dimensions**. According to the convention, the tilt axis in the raw movie frames is approximately aligned to the image X axis, while that in the tomogram is aligned to the Y axis. Thus, taking for example data recorded on a K3 detector, the unbinned image dimensions are 6000×4000 and the tomogram dimensions should be 4000×6000 instead. Set the Z dimension with this example dataset to 1000, based on the sample thickness (leave some margin at the border of tomograms to avoid problems later during subtomogram extraction).
c. Leave all preprocessing parameters/settings as those for processing of raw movie frames. Go to the **Fourier Space** tab, and click on “PROCESS ONLY THIS ITEM’S CTF” and then “CHECK TILT HANDEDNESS” to check if the defocus handedness is correct. This relates to the defocus convention and improper handling will limit the maximum attainable resolution of the subtomogram averages. If the correlation value at this step is close to 1, it indicates correct defocus handedness, as in the case of the example data. If the value is close to -1, follow instructions to flip the defocus handedness. Click on **START PROCESSING** to process the CTF for all tilt series in batch (Figure 3B).
d. When the previous step is done, go to the Overview tab and click on the **RECONSTRUCT FULL TOMOGRAMS** button. A new interface pops up that allows reconstruction of tomograms with user-defined parameters, such as the pixel size (based on the binning factor required for the next step of data processing, here 6.7752 Å used to obtain 4× binned tomograms). “Normalize input images” is useful when data are processed in RELION^11^ in the next step. Use the “Also produce deconvolved version” option with the default parameters, which significantly improving the reconstruction contrast in the example data (Figure 3C). If denoising is needed to obtain even better contrast, use the “Reconstruct half-maps for denoising” option. One can further denoise the tomograms either with Warp or other software such as cryoCARE^15^. For this data, use the “Include items outside of filter ranges” at this point. Click on **RECONSTRUCT** to start and the output tomograms will be written to the subfolder named “reconstruction” or “reconstruction/deconv” (for the deconvolved tomograms).

**Note:** Reconstruction of full tomograms at lower binning can result in slow processing, or even crashing of Warp. Also, the lower contrast in lower binning tomograms will not be beneficial for the later step of filament tracing.

Tomograms reconstructed in IMOD or AreTomo can also be used for filament tracing. But pay attention to possible pixel shifts or image rotation/flipping.

### Filament tracing and subtomogram generation

#### Timing: 6 – 9 hours

Use the filament model function in Dynamo^8^ to manually trace the backbone of the isolated filaments in the 4× binned tomograms reconstructed in Warp, generate positions for subtomograms based on the geometry of the filaments and crop subtomograms for the next steps of alignment and averaging. Our protocol was produced with an earlier version of Dynamo (1.1.401-foss-2017b-MCR-R2018a) together with MATLAB (2019b), but should be applicable to newer versions in most cases. We strongly recommend to read more detailed instructions on usage of Dynamo in the software wiki (https://wiki.dynamo.biozentrum.unibas.ch).

5. Create filament models in Dynamo.

a. Activate dynamo in MATLAB with this command: `>run /path-to-the-installation-folder-of-` `dynamo/dynamo/1.1.401-foss-2017b-MCR-` `R2018a/dynamo_activate.m` Create and enter a new working directory to run the dynamo project, here a folder named “4bin_dynamo” as an example.
b. Create a catalogue for 20 tomograms from this example data. In a terminal window (outside of MATLAB), run the command: `> ls -d /path-to-the-Warp-preprocessing-` `folder/frames/reconstruction/deconv/*.mrc >>` `list_4b_deconv.vll` This generates a text file recording the path and names of all deconvolved tomograms generated in previous steps by Warp. Then create a Dynamo catalogue named “filaments” by running: `> dcm -create filaments -fromvll list_4b_deconv.vll`
c. Generate filament models in the Dynamo graphical user interface (GUI). Open the Dynamo GUI with MATLAB command: `> dcm filaments` In the dcm GUI, click on **Create binned versions** under the **Catalogue** menu to pre-bin all tomograms with a factor of 2 again for easier visualization (target voxel size: 6.78×2^2^=27.10 Å) and filament tracing (will be annotated with the original 4× binned pixel size 6.78 Å by default in the next step by Dynamo). Then select the first tomogram at its **index** field and right click to open the binned tomogram in **tomoslice**. Follow the tutorial video (https://www.youtube.com/watch?v=hsZ-9xAYufI) to generate filament model. In Brief, call the **dtmslice** interface from the **dpreview** GUI and manually mark the two ends (“anchors”) of each filament with the [1] and [2] keyboard keys. Then create orthogonal sections along the line connecting each pair of two anchor points and click on the centers of these sections to trace the “backbones” of filaments (backbones are smoothed in later steps). If needed, modify the step parameters in the **Traverse path** box (**Sections along path** in later Dynamo versions) in the GUI to adjust how many slices to generate along the traverse path (the default 1:10:100 means to divide the distance between two anchor points by 100 parts and create a section ever 10 parts). For the example data, only modify the **sidelength** for generating the sections to 80 pixels (approximately twice larger than the filament diameter; pixel size: 6.78 Å), a value large enough to fully cover the cross-section view of sometimes curved filaments (try to avoid including heavily curved regions at the tracing step already). In the slices view, use the [C] keyboard key to choose the center of the cross-section views on different slices. Next, convert these manually clicked points into coordinates for subtomogram/particle extraction. In this case, we chose the filamentWithTorsion model (Figure 4A) to generate equidistant (dz: 8 pixels = 54.24 Å) segments of filaments with “artificial” torsion angle (dpsi: 36 degrees), in order to reduce the dominance of the missing wedge during alignment later. Finally, save the models with the positions and angles for cropping subtomograms.

**Figure 4.**
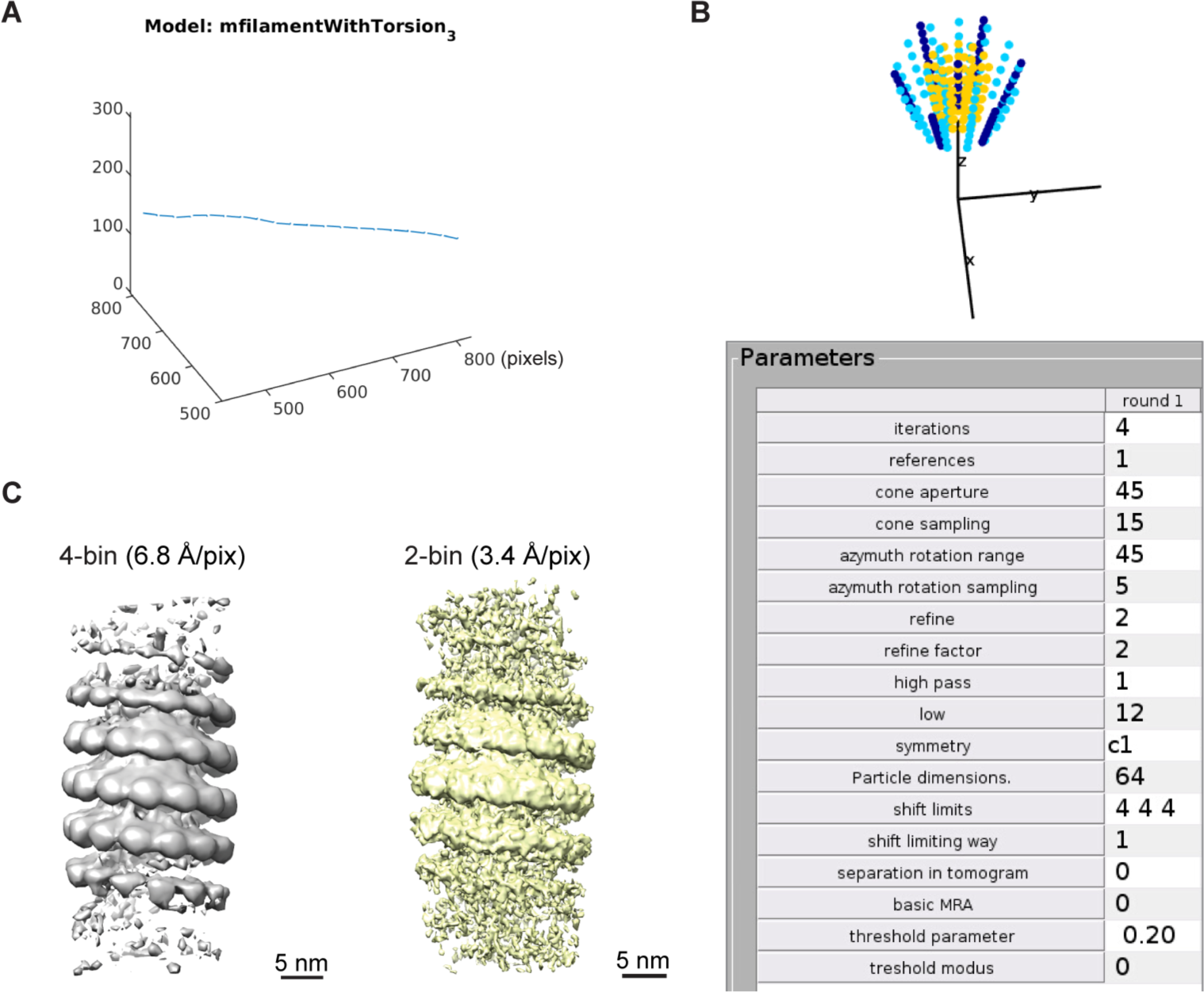
Filament model generation and initial subtomogram alignment in Dynamo. (A) One example of “filament model with torsion” generated in Dynamo. Lines with shorter perpendicular lines at their ends indicate where and with what orientations subtomograms are generated (positions and angles are recorded in a Dynamo table). (B) An illustration of the multilevel refinement of subtomogram orientations (top: dark blue, light blue, and yellow are colored according to the order of sampling grid in three consecutive levels). The corresponding numerical parameters used in the first Dynamo alignment project are shown at the bottom. (C) Averages after alignment at 4x binning and 2x binning of 1297 particles from all filaments in the example data, Y flipped to the correct handedness. Pixel sizes are indicated.

**Optional:** By incorporating loop scripting, it is also possible to input into Dynamo multiple series of points (along the traced filaments) as a matrix of their x/y/z coordinates created elsewhere (e.g. Amira) and generate geometrically computed sampling points for subtomogram extraction with MATLAB code following the Dynamo wiki (https://wiki.dynamo.biozentrum.unibas.ch/w/index.php/Filament_types_code_example).

6. Grep Dynamo tables and crop subtomograms from 4× binned tomograms with Dynamo commands in MATLAB.

a. Follow the Dynamo wiki, and use the following command lines together with loop scripting to combine multiple model tables for each tomogram into one table per tomogram.

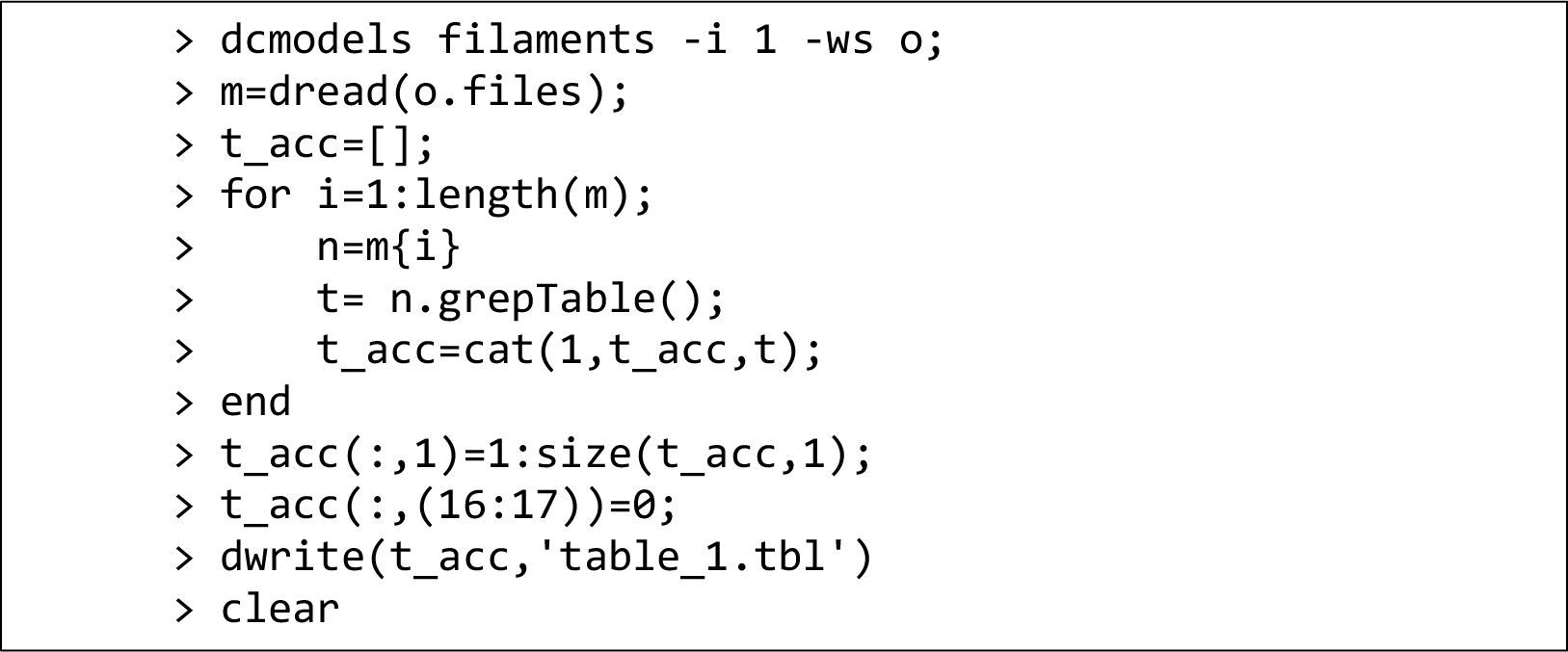 Here, the **1** after **-i** refers to the tomogram with index = 1 (also the first tomogram, explained in next steps).
b. In order to crop particles in Dynamo for all tomograms together, create a new volume list for cropping particles from the non-deconvolved tomograms in a terminal. `> ls -d /path-to-the-Warp-preprocessing-` `folder/frames/reconstruction/*.mrc >> list_4b.vll` Modify this list file so that each tomogram gets an index number and the absolute path and name of each table per tomogram (1 to 20 in this case) in this format:

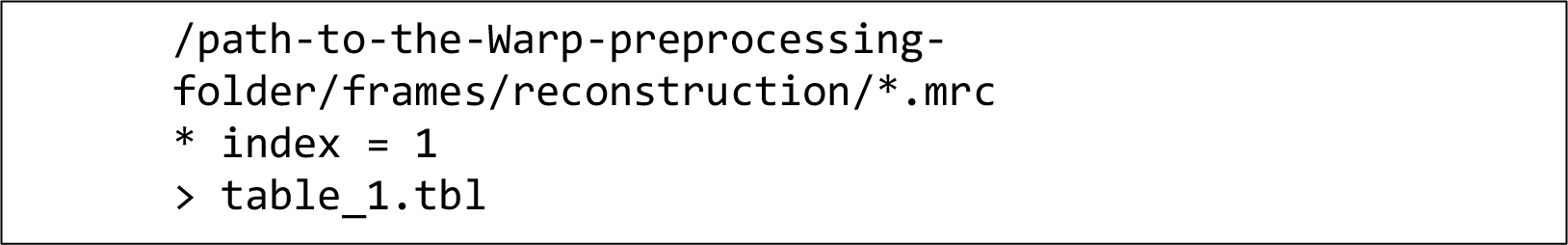 Crop all particles in MATLAB, `> dtcrop list_4b.vll reorder filamentsData 64` Here, **reorder** re-indexes particles after cropping because some positions too close to borders of tomograms will be skipped automatically by Dynamo. Also, a new table with final indices corresponding to the cropped particles is included in the particle directory. One can split the table based on tomogram indices later in MATLAB if needed. 64 refers to the box sidelength in 4× binned pixels (6.78 Å/pix), which is about twice larger than the filament diameter determined from measurements in the tomograms. Use this newly generated table (here renamed to: “table_combined_reindex.tbl”) for Dynamo subtomogram averaging in next steps. 1297 particles were generated in this step.

**Optional:** Subtomograms can also be reconstructed in Warp. Use the dynamo2warp function in the dynamo2m package to convert a dynamo table to a STAR file for Warp reconstruction (detailed later). In our initial pipeline, we converted a Dynamo table to a motl file (for TOM/AV3^9^,^10^ alignment) and/or a STAR file (for Warp/RELION) with MATLAB scripts in order to test alignment with different software at different steps.

**Note**: If MATLAB is not available, most Dynamo commands can also be run directly in the terminal as a standalone version: with Dynamo loaded as a module or path defined, one can type “dynamo” in terminal to activate it and then run Dynamo functions in a slightly modified syntax (see Dynamo wiki for more details).

There are four filament model types available in Dynamo which can be selected to fit the type of filaments being processed: filamentWithTorsion, filamenSubunitsOnHelix, filamentSubunitsOnRings, or filament (with randomly distributed positions in the filament tube).

### Initial subtomogram alignment and averaging

#### Timing: 2 – 3 days

While later high-resolution processing is performed in RELION, we found it beneficial to perform initial alignments especially for non-globular structures in subtomogram averaging-dedicated software, such as Dynamo. Align the 4× binned and 2× binned subtomograms (6.78 Å/pix and 3.39 Å/pix, respectively) with Dynamo for easy plots of aligned positions/angles and for ensuring uniform directionality of subtomograms positioned along the same filament. For detailed explanation of the parameters used in Dynamo alignment, please refer to the Dynamo wiki mentioned above.

7. Align the 4× binned subtomograms (6.78 Å/pix) in two projects (“bin4_align_1” and “bin4_align_2” in the following), mainly varying the cone and in-plane angular search parameters (Figure 4B).

a. Create an initial average of all subtomograms generated from the previous step as a template for Dynamo subtomogram alignment. With Dynamo loaded in a terminal, run the following in the working directory.

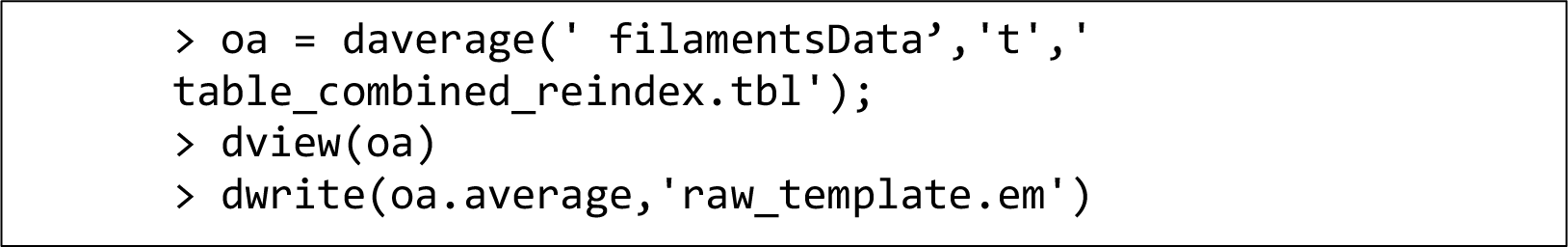
b. Set up a Dynamo subtomogram alignment project with the following command: `>dcp.new(’bin4_align_1’,’d’,’filamentsData’,’template’,’r` `aw_template.em’,’masks’,’default’,’t’,’table_combined_rei` `ndex.tbl’);`
c. Modify the alignment parameters including the masks in the GUI and run the Dynamo project. `> dcp bin4_align_1` Open the masks interface. Use the **Mask editor** to create a file, e.g. mask64_r17_h25_g2.em, with the “raw_template.em” as a reference. This corresponds to a cylinder mask with a box sidelength of 64 pixels, radius of 17 pixels, height of 25 pixels and Gaussian smoothened by a factor of 2; it should fit the template size in the overlaid view. Use this mask for the alignment, classification as well as calculating the Fourier Shell Correlation (FSC). The classification and FSC masks are not used here, but must be set for the project to run. Set the Fourier mask on template as “fill with ones” by clicking on that button at the “Input masks for project” interface. Set the alignment **numerical parameters** (Figure 4B). Key parameters are iterations: 4; cone aperture and sampling: 45 and 15 degrees; azymuth13 rotation range and sampling: 45 and 5 degrees; refine and refine factor: 2 and 2; high and low pass: 1 and 12 Fourier pixels which correspond to a maximum achievable resolution of 36 Å at the first iteration of alignment (64 (box_size) / 12 (low_pass) × 6.78 (pixel_size) = 36 Å); shift limits: [4, 4, 4] in pixels; shift limiting way: 1. Set the **computing environment** depending your local settings for running dynamo. The **Standalone** option with a few CPU cores or **GPU standalone** with the GPU identifiers properly set worked in our case. Click on the **check** and then the **unfold** button to finish preparation of the project. Exit the GUI and submit the job either by running ./bin4_align_1.exe in the Dynamo prompt, or submit the job to a cluster following the detailed instructions on the Dynamo wiki.
8. Check the consistency in the directionality of each filament, flip the ones with opposite directions, regenerate a new combined table and align the subtomograms again with this new table.

a. Split the refined table from the project “bin4_align_1” into tables for individual tomograms/filaments by reading the table into a variable and performing matrix operation in MATLAB based on the indices of tomograms in Column 20 of the table, or with the dynamo_table_grep function. Then generate averages for individual tomograms from the split tables with the daverage and dwrite functions as shown above. One can automate and combine these two steps with a loop script in MATLAB.
b. Inspect the averages for individual tomograms in the previous step and takes notes of the ones showing opposite directions to the first subtomogram average (used as a reference here). In our case, we found that Tomogram 1, 2, 7, 10, 13 gave averages of opposite directions (single filament in each tomogram). Flip the directions for these subtomograms by operating on their corresponding tables and adding 180 degrees to the values in Column 8 of these sub-tables in the version before the execution of any alignment, with the following MATLAB script.

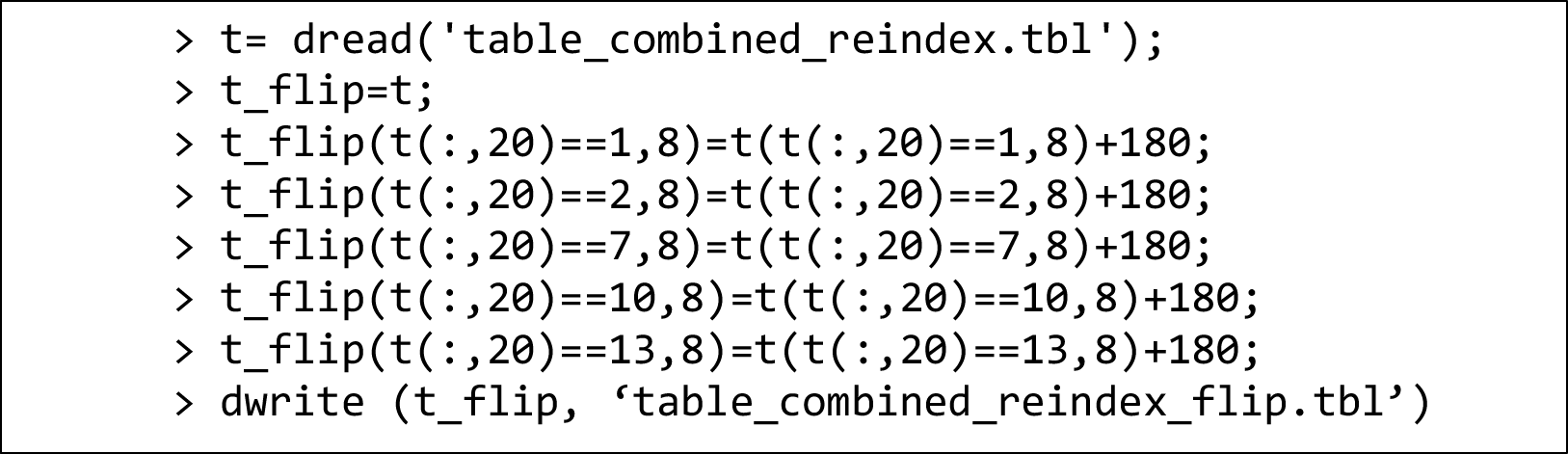
c. Run a new alignment project named “bin4_align_2”. Set the masks as for previous alignment projects and the alignment parameters as the following: cone 60/15, in-plane 45/15, refine 2/2, high/low pass 1/12, iter 4, shift limits [4, 4, 4], shift_limiting way 1.
9. Run a Dynamo adaptive band pass (ABP) filtering project from the refinement results of “bin4_align_2”.

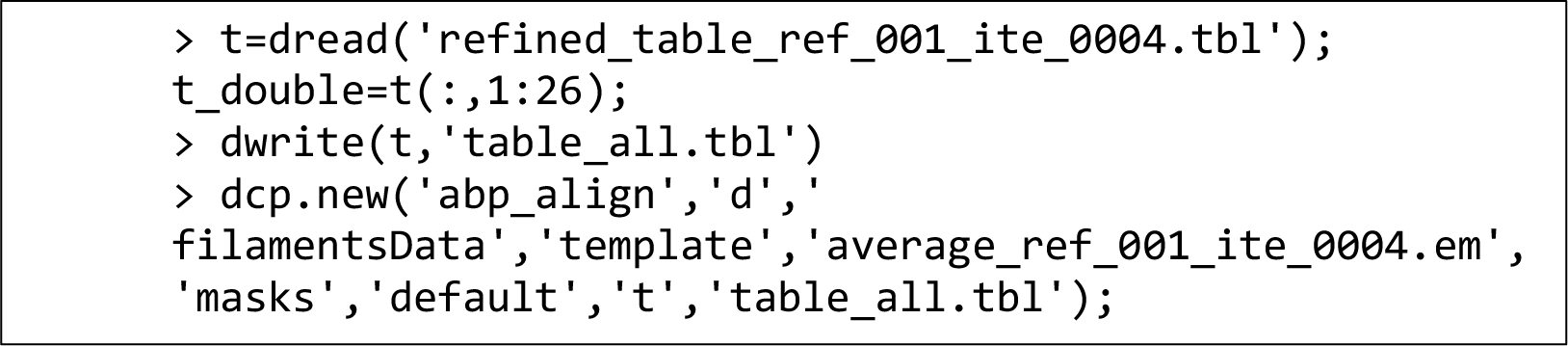 Launch the project GUI with the dcp command and set the same masks as for previous alignment projects. Now, the FSC mask will be meaningfully applied to each of the half averages at each iteration for estimating resolution and defining the lowpass filter parameter for the next iteration. Then derive an ABP project from this single reference project: in the GUI choose the following options in the **Multireference** tab > **Adaptive filtering (“golden standard”)** > **Derive a project**. In the newly launched interface, rename the derived ABP project as “bin4_align_2_eo” following the convention and click the Enter key. Generate the odd/even references by averaging the odd/even subtomograms in the eo project GUI (at the template interface). Also, at the template interface, use the **visualize FSC** button to estimate the initial frequency given in Fourier pixels (the X value corresponding to a Y value of around 0.143 in the FSC plot when moving the cursor along the curve^16^) that will be used as the lowpass filter parameter of the first round of alignment. Set this X value together with an FSC threshold of 0.143 and a resolution pushback of 2 Fourier pixels in the dcp GUI under the **Multireference** tab > **Adaptive filtering (“golden standard”)** > **Edit parameters for Adaptive Filtering run**. Then set the numerical parameters and computing environment as for previous Dynamo projects. Run the project. When finished, launch the project GUI again and use the **Plot attained resolution FSC for all iterations** option under the **Adaptive filtering (“golden standard”)** option to check the evolution of FSC plots and resolutions in Fourier pixels in the different iterations. This project converged and resulted in an average (Figure 4C, left) with an estimated resolution of 64/19*6.78=22.8 Å (1 / Fourier_pixels * box_size * pixel_size; 19 Fourier pixels here).
10. Combine the tables from the bin4_align_2_eo project and prepare for subtomogram reconstruction and alignment at 2× binning (pixel size: 3.39 Å). Due to the difficulty and unnecessity in reconstructing full tomograms at 2× binning for Dynamo reconstruction, we choose Warp for reconstructing subtomograms at lower binning. In the Warp GUI at the .tomostar mode, click on the **RECONSTRUCT SUB-TOMOGRAMS** button to create subtomograms and their corresponding 3D CTF models (models and the output STAR file are not used at 2× binning but will be important for the next step processing of the unbinned subtomograms). In the pop-up options for exporting subtomograms, set the “Coordinates use 6.7752 Å/px” based on the coordinate scale in the STAR file converted from a Dynamo table; set the “Output will be scaled to 3.3876 Å/px” for 2× binned subtomograms; set the “Box size is 128 px”; set the “Particle diameter is 433 Å” with 433 as the integer of 3.3876×128=433.6128 to include most signal/noise in the box; select the following options “Volumes”, “Invert contrast”, “Normalize input images”, “Normalize output volumes”, “Make paths relative to STAR”, and “Include items outside of filter range”. Rename the subtomogram files from the Warp convention (e.g. f00003_0000001_3.39A.mrc) to the Dynamo convention (e.g. particle_000100.em) and convert the mrc files to em files for Dynamo. Multiple methods such as the the tom_mrcread and tom_emwrite functions in the TOM package^9^ within MATLAB, the mrc2em module in PyTom^17,18^, or the e2proc3d.py in EMAN2^19^ can be used to perform this conversion.
11. Align the 2× binned subtomograms by creating a new Dynamo ABP project, following the same procedures described above (generate a new mask of twice the size of that used for 4× binning; decide on the initial lowpass frequency by calculating from the new odd/even averages at 2× binning; use smaller angular and shift search ranges). When the ABP project at 2× binning (pixel size: 3.39 Å) converges, i.e. reaching a resolution of 13 Å in this case (Figure 4C, right), prepare for alignment of unbinned subtomograms (pixel size: 1.69 Å) in next steps. First, clean up clashing subtomograms (particles that have shifted to occupy the same position in the helical filament) based on their distances to each other and their cross-correlation (cc) values within MATLAB.^20^ Here, a low threshold (10 pixels at 2× binning, i.e. 3.39 nm) was used after checking the histogram of distances of subtomograms to their 20 nearest neighbors. For the clashing subtomograms, the ones with higher cc values were retained. Here, 1209 subtomograms were kept for the next step processing. Convert the cleaned Dynamo table to a STAR file for Warp and reconstruct the unbinned subtomograms in Warp. Perform one round of Dynamo alignment to make sure that the cleaned table/STAR file works before proceeding to RELION in the next steps.

**Note**: One can also use the average of subtomograms from a single long filament as an initial template. As we have tried to keep uniform directionality during the filament tracing step, the average of all subtomograms gives better signal-to-noise ratio and should not be dominated by subtomograms from filaments with opposite directionality.

Dynamo alignment parameters can also be set simply by command lines without the GUI.

For initial alignments at 4× binning, one can also try using a large cone for angular search (e.g. 360/45). With the example data, such test resulted in a worse average showing fake 2-fold symmetry. Thus, for later steps of alignment at lower binning, even smaller angular and shift limits than the first alignment is recommended.

Use dynamo_table_plot function to plot orientation and positions of subtomograms after refinement. Dynamo has multiple flags regarding how the plots are displayed, e.g. colored according to cc values. The values in tables can be plotted in histograms, e.g. the shifts in the refined tables, which helps for evaluation and optimization of the refinement procedures beyond visualization of the resulting averages.

Originally, we aligned the 2× binned subtomograms in TOM/AV3 instead of Dynamo for speeding up the alignment in parallelized jobs essentially on the cluster (not working efficiently in Dynamo versions installed on our cluster with the corresponding setup).

### Subtomogram alignment and helical refinement with unbinned data

#### Timing: 11 days

Align the unbinned subtomograms (pixel size: 1.6938 Å) with refined positions and orientations from the steps above in RELION. Perform 3D auto-refine, helical search, 3D auto-refine with helical reconstruction, and 3D classification (Figure 5; RELION version: 3.1.0). We recommend referring to the guidelines for RELION (https://www3.mrc-lmb.cam.ac.uk/relion/index.php/Helical_processing) and its helical processing instructions for more details about RELION conventions (https://www3.mrc-lmb.cam.ac.uk/relion/index.php/Refine_a_structure_to_high-resolution).

12. Run RELION 3D auto-refine without helical reconstruction.

a. The Warp-generated STAR file from the last step is in a format compatible with earlier RELION versions. For RELION version 3.1 or later, use the relion_convert_star command (with the RELION module loaded) to convert an old STAR file format to a new one whenever needed. **<u>Troubleshooting 3</u>**

**Figure 5.**
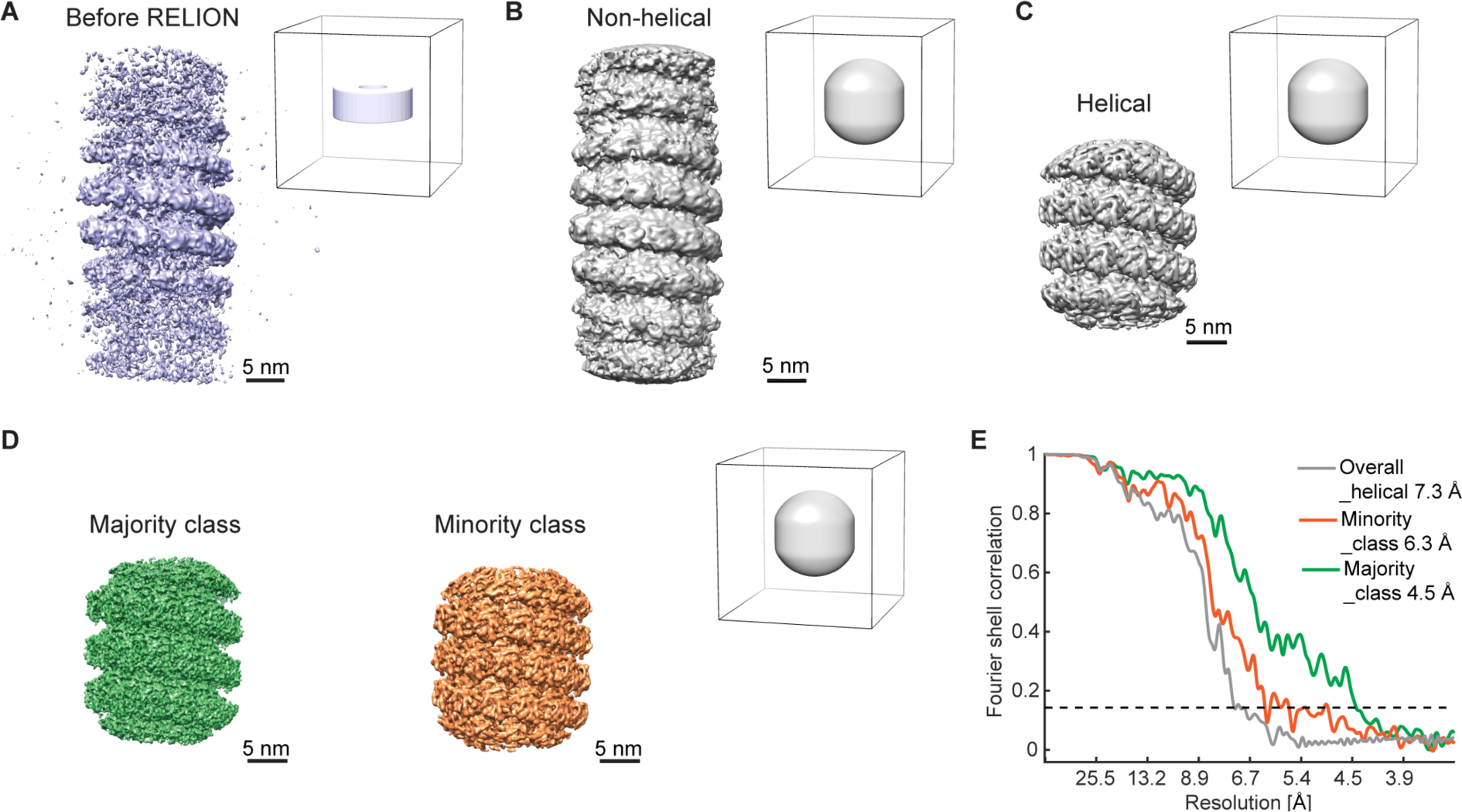
Subtomogram averages imported into RELION for refinement and helical reconstruction. Pixel size is 1.69 Å (unbinned data) for all averages. (A) and (B): half map, Gaussian filtered, without multiplication by the alignment mask (shown next to each map). (C) and (D): half map, filtered to the determined resolutions in RELION Post-processing jobs, with the alignment mask (next to the map) multiplied. The maps in (D) are final results after classification for each class following M refinement. All maps are Y flipped to the correct handedness. (E): masked FSC curves for the maps in (C) and (D), using the same alignment mask. The dotted line indicates the 0.143 threshold.

**Note:** New-style RELION STAR files (version 3.1 onwards) mainly differs from old ones in: 1) the new-style includes the optics group part; 2) the new-style uses OriginX/Y/Zangst columns to replace the OriginX/Y/Z and the units change from pixels to angstrom. Thus, these values in the new STAR files equal values in OriginX/Y/Z × pixelSize in angstrom. We used a MATLAB script for this conversion. The dynamo2m package enables this with python scripting.

b. Convert the average of the last round of Dynamo alignments from a .em file to a .mrc file in Chimera^12^. Use the Volume Viewer/Tools/Volume Filter to scale the volume by -1. As the Dynamo average likely does not contain pixel size information, set the correct pixel size (1.6938 Å) with Chimera under Volume Viewer/Features/Coordinates. Then save the map as a new map named ref_unbin.mrc (Figure 5A).
c. Generate a cylinder mask with RELION. Use the command as following: `>relion_helix_toolbox --cylinder –o` `cylmask256_o220_s55_w5.mrc --boxdim 256 –` `cyl_outer_diameter 220 --angpix 1.6938 --` `sphere_percentage 0.55 --width 5` Check the RELION instructions for more explanations about these parameters. In brief, measure the filament outer diameter based on ref_unbin.mrc. The combination of boxdim (in pixels), cyl_outer_diameter (in Å), sphere_percentage gives a central Z length of about 30%, which is optimal for RELION helical reconstruction in later steps. Width (in pixels) refers to the value for creating a soft-edged mask.
d. Set up the RELION job in a GUI. Leave the setup parameters as default unless specified otherwise. Set the newly converted STAR file, the ref.mrc, and the cylinder mask file in the I/O tab. In the Reference tab, leave the “**Ref. map is on absolute greyscale?**” as No because the map was not generated in RELION and thus may have a different absolute intensity grey-scale from the RELION requirement. Set the **Initial low-pass filter** as 30 Å. In the Optimisation tab, set the **Mask** diameter as 300 Å and the “**Use solvent-flattened FSC?**” as Yes. In the Auto-sampling tab, set the **Initial angular sampling** as 3.7 degrees and the **Local searches from auto-sampling** as 3.7 degrees. For the Compute and Running tab, set parameters according to the computing environment. In our case, we copied the particles and their corresponding 3D CTF models to a scratch directory on a cluster and set the **Number of MPI procs**, **Number of threads**, **GPU count**, **Memory**, **Walltime** and **Minimum dedicated cores per node** as 3, 7, 2, 120GB, 3-23:00:00, 15, respectively. Click on **Run** to submit the RELION job to a cluster. It took 33 iterations to converge in this case.
e. In RELION Post-processing, provide **One of the 2 unfiltered half-maps** and the refinement mask as the **Solvent mask** and provide a B-factor of for example -500. When the job is done, check the estimated final resolution in the output window, as well as in the output files. The resulting resolution was 10 Å with the refinement mask applied, showing clear separation of subunits along the helical turns (Figure 5B).

13. Perform helical parameter search in RELION. With the output average from the Post-processing job in Step 12, estimate the helical rise and twist parameters by measuring the helical pitch and counting the number of subunits per turn (rise about equals the helical pitch in Å divided by the number of subunits per turn; twist about equals 360 degrees divided by the number of subunits per turn) in order to set the ranges in the following command line: `>relion_helix_toolbox --search --i xxx.mrc --cyl_outer_diameter` `220 --angpix 1.6938 --rise_min 3.5 --rise_max 4.5 --twist_min` `25 --twist_max 35 --z_percentage 0.3` The output from this command is: rise 4.46 Ang; twist 27.08 degree.
14. Run RELION 3D auto-refine with helical reconstruction. The settings are similar to those in Step 12, except for the following:

a. Use the output STAR file from the 3D auto-refine job in Step 12.
b. Use the output average from the Post-processing job in Step 12 as reference map.
c. Set up the RELION job in a GUI. Set the STAR file, the reference map, and the reference mask file (the same as in Step 12) in the I/O tab. In the Reference tab, leave the “**Ref. map is on absolute greyscale?**” as Yes. In the Helix tab, change the following parameters: ***Do helical reconstruction?** Yes;* ***Tube diameter – inner, outer (Å)**: -1, 220;* ***Keep tilt-prior fixed**: Yes;* ***Number of unique asymmetrical units**: 13*; ***Initial twist (deg), rise (Å)**: 27.08, 4.46;* ***Do local searches of symmetry?** Yes;* ***Twist search – Min, Max, Step (deg):** 25, 29, 1* ***Rise search – Min, Max, Step (Å)**: 3.8, 4.6, 0.05* In the Running tab, increase the **Walltime** as this job took longer than the refinement job without helical reconstruction with the RELION version used. Click on **Run** to submit the RELION job to a cluster.
d. In RELION Post-processing, use one of the two unfiltered half-maps of the previous step and the refinement mask as the Solvent mask and perform B-factor estimation automatically. The resulting resolution was 7.3 Å with an “Auto b-factor” of -382. Re-doing helical search gives a helical parameter set of 27.16° and 4.17 Å (Figure 5C).
15. Run RELION 3D classification with helical reconstruction to check for heterogeneity in the particles. Settings are left as default, except for the changes in the following:

a. Use the output STAR file from the 3D auto-refine job in Step 14.
b. Use the output average from the Post-processing job in Step 14 as the reference map.
c. Set up the RELION job in the GUI. Set the STAR file, reference map, and the cylinder reference mask file (the same as in Step 12) in the I/O tab. In the Reference tab, leave the “**Ref. map is on absolute greyscale?**” as Yes. Set the **Initial low-pass filter** as 40 Å. In the Optimisation tab, set the **Number of classes** as 4, the **Mask** diameter as 300 Å. In the Sampling tab, set the **Angular sampling interval** as 1.8 degrees, the “**Perform local angular searches**?” to Yes and use a **Local angular search range** of 10 degrees. In the Helix tab, change the following parameters: ***Do helical reconstruction?** Yes;* ***Tube diameter – inner, outer (Å)**: -1, 220;* ***Keep tilt-prior fixed**: No;* ***Number of unique asymmetrical units**: 13;* ***Initial twist (deg), rise (Å)**: 27.16 and 4.17;* ***Do local searches of symmetry?** Yes;* ***Twist search – Min, Max, Step (deg):** 26, 28, 1* ***Rise search – Min, Max, Step (Å)**: 3.3, 4.5, 0.05*
d. The result has 4 classes, of which Class 3 contained 77.7% of the particles (939 particles) and had a resolution of 5.9 Å, refined helical twist = 27.1713°, rise = 4.21176 Å. Class 4 contained 21.3% of the particles (258 particles) and had a resolution of 6.6 Å, refined helical twist = 26.9342°, rise = 3.51426 Å. The other two classes containing 12 particles in total did not result in any meaningful average.
16. Run RELION 3D auto-refine with helical reconstruction for Classes 3 and 4 separately. Settings are similar to those in Step 14, except for changes in the following:

a. Use the selected STAR file for particles in Class 3 from the classification job, and that for Class 4 in a separate job.
b. Use the output average from the Classification job as Reference map (average from Class 3 and 4 separately for two jobs).
c. Set up the RELION job in a GUI. Set the STAR file, the Reference map, and the cylinder Reference mask file (the same as in Step 12) in the I/O tab. In the Reference tab, leave the “**Ref. map is on absolute greyscale?**” as Yes. Set the **Initial low-pass filter** as 20 Å. In the Helix tab, change the following parameters: ***Do helical reconstruction?** Yes;* ***Tube diameter – inner, outer (Å)**: -1, 220;* ***Keep tilt-prior fixed**: Yes in principle; if the RELION job aborts with the error ”Segmentation fault”, set this to No and re-run it*. ***Number of unique asymmetrical units**: 13;* ***Initial twist (deg), rise (Å)**: 27.17, 4.21 for Class 3 and 26.93, 3.51 for Class 4;* ***Do local searches of symmetry?** Yes*; ***Twist search – Min, Max, Step (deg):** 25, 29, 1* ***Rise search – Min, Max, Step (Å)**: 3.8, 4.4, 0.05*
d. In RELION Post-processing, the resulting resolution for Class 3 was 6.5 Å with an Auto b-factor of -85. Re-doing helical search for the Class 3 average gives a helical parameter set of 27.17° and 4.21 Å. The resolution for Class 4 is 7.3 Å with an Auto b-factor of -161. Re-doing helical search for the Class 4 average gives a helical parameter set of 26.91° and 3.51 Å (Final refinement maps of these two classes are shown in Figure 5D with corresponding FSC curves in Figure 5E).

**Note**: RELION had difficulty in maintaining the angles imported from initial refinement steps in other software.

EM maps at a resolution of 8 Å or higher should exhibit some features of secondary structures such as helices, which should be critically assessed along the alignment/refinement steps, rather than the nominal estimated resolution.

### Per-particle refinement and map evaluation

#### Timing: 3 days

Perform per-particle refinement in M (following helical symmetry expansion with RELION, as the M version until 1.0.9 does not support helical symmetry operation) and rerun RELION 3D auto-refine with refinement results from M. Generate FSC plots and examine local resolution of maps.

17. Prepare files for M refinement.

a. Make a copy of the original Warp preprocessing, especially the .xml and .tomostar files as a backup. M refinement will overwrite some of the files.
b. Copy the following files (2 half maps and particle STAR file) from a RELION 3D auto-refine job to a folder designated for M processing (ideally as a subfolder in the directory that contains the Warp frames folder): it0xx_half1_class001_unfil.mrc, it0xx_half2_class001_unfil.mrc, and run_it0xx_data.star (renamed to run_it0xx_data_ori.mrc)
c. For an M version that does not support helical symmetry operation, perform helical symmetry expansion with the RELION command: `>relion_particle_symmetry_expand --i run_it0xx_data.star` `--o run_it0xx_data_expanded.star --helix true --twist` `27.168 --rise 4.21387 --angpix 1.6938 --asu 13` This generates a new STAR file with an expanded particle list according to the symmetry (13-fold larger). Downgrade the RELION STAR file with the relion_star_downgrade command in the dynamo2m toolbox and rename the STAR file to jobxxx_run_it0xx_data.star for M refinement (M version 1.0.9 needs to recognize the suffix of the file as _data.star).
d. Copy the cylinder mask that was used for RELION refinement to the M processing folder.
18. Run M refinement.

a. Open the Warp session before running M refinement to make sure the correct settings/input files can be imported by M. Close Warp and open M. Create a new population in the working directory that contains the Warp frames folder. Link the new population to the data source by clicking “Manage data sources”, then “ADD LOCAL” at the interface. Choose and import the “previous.settings” file from the previous Warp processing folder. Set a name for the data source and a source file will be created. Choose to “Include items outside of filter ranges” to include all data selected. Continue to click “Create” and close the pop-up interface when done. A .source file will be created within the frames folder by default.
b. Click the “+” icon to add species, for example the Class 3 average obtained in step 16. Use the “From scratch” option for the first M refinement job and click on “ADD”. Set a name for the species and use a parameter set of 220 Å diameter (filament diameter), 2600 kDa (the estimated molecular weight of the region included in the refinement mask, estimated based on 39 protein subunits plus 6 × 39 = 234 nucleotides for RNA; with a small margin), C1 symmetry, 1 temporal samples for poses. Select the half-maps from the M processing folder above for the Class 3 average for example and type the correct pixel size. Use the “filter both to Å” option to visualize the input maps (filtered to a resolution lower than the estimated value, for inspection purpose). Set the mask file with the cylinder mask in the M processing folder. Set the symmetry expanded and downgraded STAR file. Input the pixel size for coordinates and shifts in the STAR file which relates to the binning factor at which alignments are performed. Select the source file under “Use these sources for name matching”. M then displays the correct particle number read from the STAR file that can be matched to this data source file. All particles should be found if the input STAR file format is correct and the source file contains all corresponding tomograms. Click on “FINISH” when done with these input settings. M starts preprocessing steps, including denoising, which may take 10 minutes. A GPU memory issue may cause software crash at this step (GPU memory of about 24 GB or above is recommended).
c. M reports an initial resolution comparable to that obtained from RELION auto-refinement job. Click on the “REFINE” button on the upper right corner in the M GUI, to set/enable the following refinement settings in the first round: *Refine for 3 sub-iterations, use 70% of available resolution in first sub-iteration* *Image warp grid: 4 × 4* *Particle poses* *Stage angles* *Volume warp grid: 3 × 3 × 2 × 10* This provides an improvement in resolution from 6.5 Å to 6.1 Å for Class 3 as an example. Continue with a number of additional rounds of M refinement until the resolution stops improving. Add the following refinement options for a second stage of M refinement: *Anisotropic magnification* *Tilt movies* *Use species with at least 7.0 Å resolution* *Defocus and grid search in 1^st^ iteration* Continue with a few additional rounds of M refinement until the resolution stops improving.
d. When the refinement is done, M gives an estimated resolution of 4.8 Å in the refined map. Inspect the refined and postprocessed maps, as well as the corresponding FSC curves generated by M refinement in the species folder. The *fitsharp* and *denoised* maps should show improved details in the maps compared to the input map, e.g. better recognizable helices or even large side chains of residues at this resolution range. The FSC curve should drop to zero at the high frequency range and not show “bumps”. If satisfied with the quality, continue to the next step.
19. RELION refinement after M. This further improved the resolution of the EM map for the majority class from 4.8 Å to 4.5 Å.
  a. Convert the STAR file from M (_particles.star in the latest M species folder) for reconstruction of particle/subtomograms in Warp with the M-refinement-updated poses. We performed this with a MATLAB script that converts the coordinates unit from Å back to pixel, removes 12 extra positions of the symmetry-related particles per 13 particles. Modify the data columns in the STAR file using a previous STAR file for Warp as a template. Reconstruct subtomograms and their 3D CTF models with this STAR file and convert the STAR file again for RELION versions above 3.1.
  b. Prepare a new reference from the M refinement result. The M generated average (the filtered and sharpened *fitsharp* map in the species folder) does not use the same box as the subtomogram particles for RELION. Therefore, open a pervious RELION average (read as #0 in Chimera) and the M average (read as #1 in Chimera) in the same Chimera window, superpose the two averages by the “fit in map” option and use the following command to rescale the box size and set the origin for the M average: `>vop resample #1 onGrid #0` Save the resampled average in .mrc format and use it as an input reference in a new RELION 3D auto-refine job.
  c. Run a new RELION 3D auto-refine job with helical reconstruction, with settings similar to Step 16. Use a smaller auto-sampling angle as the particle orientations have been refined extensively in previous steps. Do post-processing for this new average to obtain a final refined map. Search for helical parameters again with the post-processed average. The parameters remained almost the same after M refinement compared to those before, but the calculated resolution with the 0.143 criterion and map quality improved substantially with M refinement (Figure 6).
20. Map quality evaluation and model building.

a. Use RELION modules to calculate the global and local resolutions of the final maps. As mentioned above, a RELION Post-processing job generates FSC plots based on the unfiltered half maps, with or without a given mask. The resolutions we reported as 4.5 Å for the majority class average and 6.3 Å for the minority class average were derived with the cylinder refinement mask applied. The maps after M-RELION refinement show higher quality in that secondary structures can be easily visualized by eyes in the final maps. One can then run a RELION Local resolution job to calculate the local resolution and render the map accordingly in Chimera/ChimeraX^13^.
b. Prepare for build a structural model based on the final map and check the model-FSC correlation. As mentioned above, we generated the maps with incorrect handedness using Warp 1.0.7 beta. Before building the model, first run the command line below to get the handedness of the map corrected:

**Figure 6.**
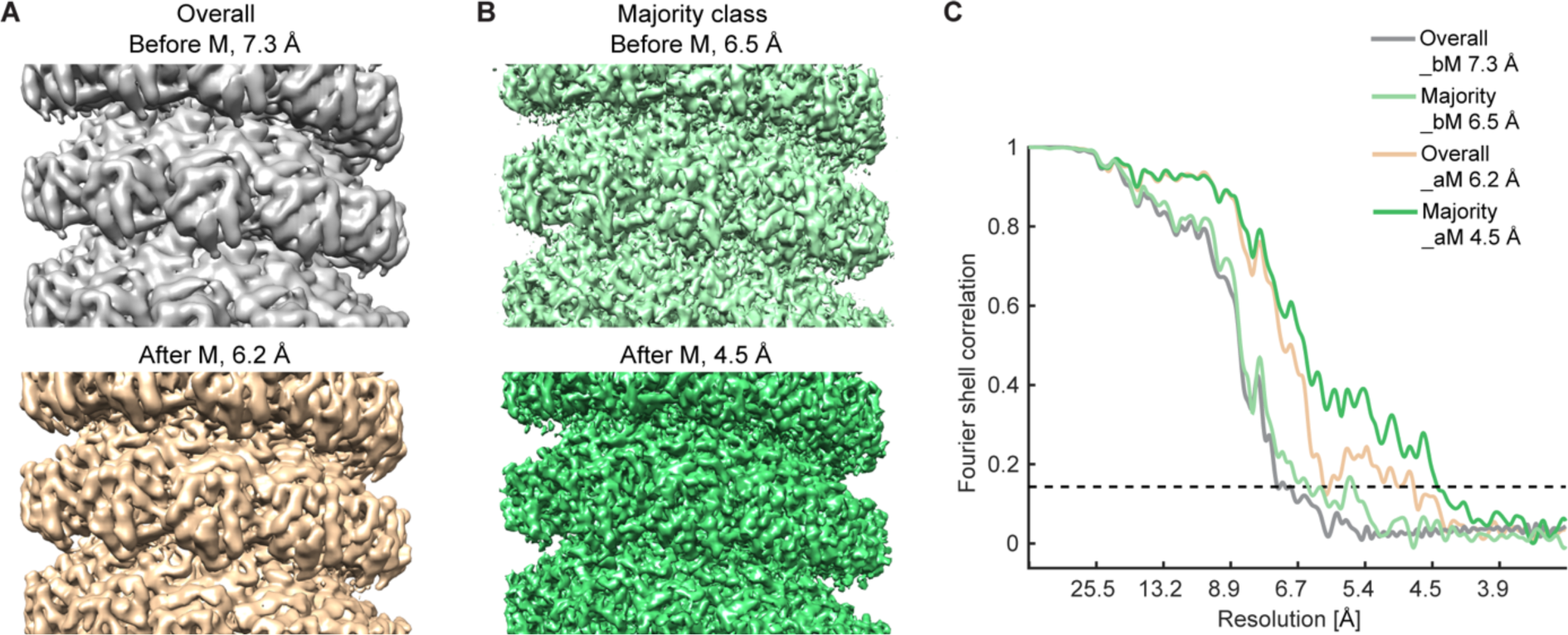
Map comparison of subtomogram averages before (top) and after (bottom) M refinement, from all the particles (A) or those belonging to the majority class (B), along with their masked FSC curves (C). The alignment mask shown in Figure 5D was used for refinement and postprocessing of all averages shown here. The dotted line in (C) indicates the 0.143 criterion. bM: before M refinement; aM: after M refinement. Numbers refer to the determined map resolution. All maps are Y flipped to the correct handedness. `>relion_image_handler --i xxx.mrc --o xxx_flipY.mrc --` `flipY` For building a model, as detailed in Zhang et al.,^1^ we generated an initial homology model with I-TASSER^21^, before AlphaFold^22^ became available. The I-TASSER model served as a starting point for iterative automated and manual refinement in Phenix^23^, and *Coot*^24^. A tetramer model was built in order to better analyze the interaction interface among monomers of the same turn as well as between the upper and lower turns. At the end, generate a segmented map for the region corresponding to the tetramer model with the **volume zone** command in ChimeraX and validate the real-space per-residue cross-correlation of the tetramer model to the map generated in Phenix.

**Note:** M version 1.1 beta could support helical refinement but needs be tested. According to the developer of M, one has to modify the **“Symmetry” Value** (e.g., Hel|27.17|4.21) in the .species file as setting the helical parameters is not yet supported directly in the GUI.

In case M refinement does not improve the resolution of maps, but rather deteriorates it, make a copy of all files from an earlier version of that species (contained in **[species folder]/versions**, sort by date to get the version needed) except for the [species name]_particles.star file, to get back to the maps with higher resolution and try other combinations of refinement options.

For non-helical objects, one can use the export sub-tomograms function in M instead of resorting back to Warp with a converted STAR file.

For M refinement of particles of a small number (e.g. Class 4 from Step 15), set up a refinement job with two species using STAR files and half maps for both species (Classes 3 and 4 together). Refinement worked when including only the image warp grid, particle poses, stage angles, and volume warp grid options. We continued to reconstruct subtomograms after M refinement and run a new RELION 3D auto-refine job for the results described here.

## Expected outcomes

Based on 20 cryo-tomograms, which contained about 20 viral nucleocapsids, we extracted 1297 subtomograms from the relatively straight nucleocapsid segments and successfully obtained 2 reconstructions at 4.5 Å and 6.3 Å. This 4.5 Å maps was of sufficient quality to serve as the basis for the building of an atomic model of the mumps viral nucleocapsid protein (N terminal 3-405 amino acids) with its RNA hexamer as one subunit for the assembly of the helical filamentous structure. We then used this model to interpret the 6.3 Å lower resolution minority class reconstruction by careful visual inspection in Chimera/ChimeraX, and provided information on the differences in helical pitch and assembly compactness between the two maps. Comparing the two models further revealed differences in the interaction interfaces between monomers leading to the different assemblies, as well as their conformations that indicate differences in the surface accessibility of their bound RNA (genome) and C-terminal regions (Figure 7).

**Figure 7.**
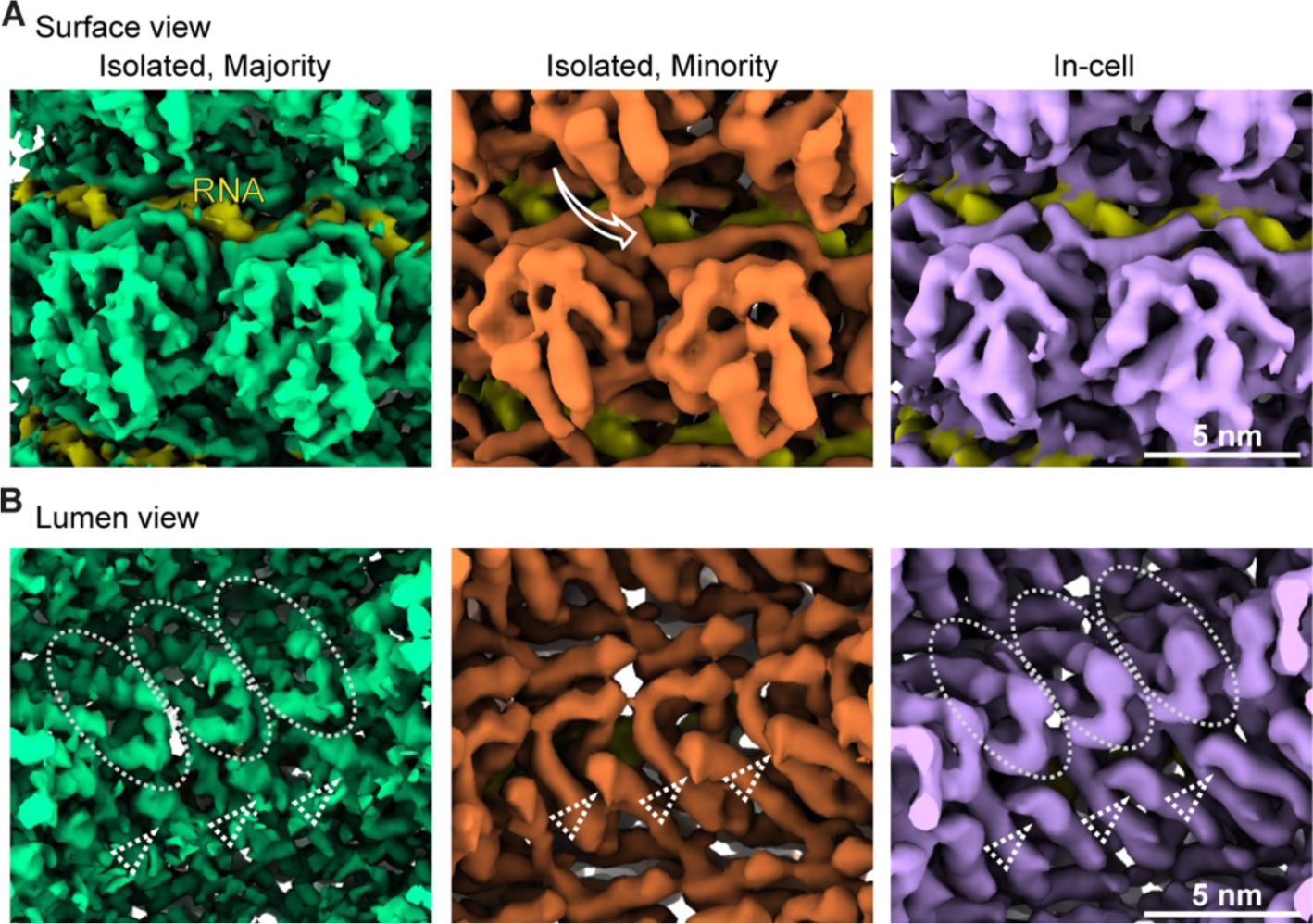
Comparison of subtomogram averages of the majority (green) and minority (orange) classes of the isolated nucleocapsids, and of the average from the in-cell nucleocapsids (purple), shown form the outer surface view (A) and lumen view (B). RNA is depicted in yellow in all images. The white arrow in (A, middle) indicates the shift of subunits in the upper turn towards the lower turn in the minority class average, in comparison to the other two averages. The white circles in (B) denote a region in the density which is not seen in the minority class average possibly due to flexibility, in contrast to density consistently seen in all averages (white arrowheads).

Beyond the example dataset of extracted helical nucleocapsids described in this protocol, we have also processed cellular cryo-ET data of relatively straight nucleocapsids using steps similar to those discussed here, with the following adjustments: 1) the tracing of the helical filaments was performed in Amira using dedicated functions^1^; 2) the generation of equidistant sampling points along the helical segments was done with a MATLAB script^25^; 3) initial alignment of the subtomograms required more strict control of the Euler angles to avoid change of directionality of subtomograms along their respective filaments; 4) checking the directionality of each filament was complicated by the lower signal-to-noise ratio of the cellular cryo-ET data compared to that of the isolated filaments. Thus, the ones with unclear directionality based on the inspection of the per-filament average were largely discarded at the initial alignment steps for achieving high resolution. Nevertheless, a consensus map of 6.5 Å was finally obtained from 5 tomograms, 68 helical filaments and 2178 subtomograms.

## Limitations

Transfer of the workflow described in this protocol to the processing of different filament types and morphologies is expected to be successful, provided that the filaments are sufficiently straight or contain straight regions. Depending on the filament types and available software in your local settings, data processing workflows that integrate other software, such as those described for the processing of LRRK2-microtubule filaments^26^, the COPII coat reconstituted on lipid membranes^2,27^, the Ebola virus nucleocapsid^28^, or the skeletal thin filaments^29^ can also be tested.

## Troubleshooting

### Problem 1

Uncertainty about the options for correcting gain reference during Warp preprocessing (related to Step 1).

### Potential solution

- The correct gain options need to be tested according to the camera setting and serialEM data saving options. Warp indicates whether the “Transpose” option is needed automatically. According to the “r/f” values of the raw tif images (https://bio3d.colorado.edu/SerialEM/hlp/html/about_camera.htm#norm_frames), different combinations of “Flip X axis” and “Flip Y axis” are needed.
- Using images with good contrast, test the flip combinations by processing a few movies without motion correction until a visible gain pattern disappears in the average images. Or, generate a sum of dozens of averages without motion correction. Check the gain pattern in this summed image to decide on the flip options.

### Problem 2

Inconsistency between Warp and IMOD reconstruction (related to Step 4).

### Potential solution

- Exclude tilt views only during IMOD reconstruction, not before. The Warp 1.0.9 version can recognize these excluded tilt views, including those excluded at the starting page of Etomo reconstruction due to largely dark regions, or later at the fine alignment step due to large residual errors. Thus, there is no need to manually edit each mdoc files for these excluded views.
- Alternatively, manually exclude tilt views in Warp before creating tilt series for IMOD, and edit the corresponding mdoc files.
- When using AreTomo or other software that performs automatic reconstruction, it is recommended to first test with one tilt series to solve possible inconsistency before batch processing.

### Problem 3

RELION refinement errors related to particle path not found or STAR file style (related to Step 12).

### Potential solution

- It is important to update the path of subtomograms and 3D CTFs in the STAR files whenever the data path or folder name is changed. Using an absolute path instead of a relative path mostly avoids problems.
- The new STAR file style poses an issue of conversion when using Warp, RELION, and M. Warp/M until the version 1.0.9 takes only the old style of STAR file, while RELION from the version 3.1 onwards uses the new style. Although RELION 3.1+ versions may automatically upgrade old-style ones, errors could still happen as “not all necessary variables defined in _optics.star file”. Pay special attention to the rlnPixelSize, rlnVoltage and rlnSphericalAberration values and manually add these lines if not found in the STAR file after using the relion_convert_star command.

## Resource availability

### Lead contact

Further information and requests for resources and reagents should be directed to and will be fulfilled by the lead contact, Julia Mahamid.

### Materials availability

- This study did not generate new unique reagents.

### Data and code availability

- Raw micrographs are publicly available in the Electron Microscopy Public Image Archive (EMPIAR) under accession code 10751 (isolated nucleocapsids). Cryo-EM maps are deposited in the Electron Microscopy Data Bank (EMDB) under accession codes EMD: 13133 (majority class, isolated) and EMD: 13136 (minority class, isolated). The associated atomic model is deposited in the Protein Data Bank PDB under accession code PDB: 7OZR.
- This paper does not report original code.
- Any additional information required to reanalyze the data reported in this paper is available from the lead contact upon request.

## Acknowledgments

We thank the EMBL EMCF, central IT and Thomas Hoffmann for computational support, Wim Hagen, Felix Weis, and the cryo-EM platform for support in data acquisition. We thank the Mahamid group members for discussions and help. We are grateful to Giulia Zanetti, Daniel Castano-Diez, and Beata Turonova for initial advice on subtomogram averaging strategies, Dimitry Tegunov for developing helical symmetry functionality in M. J.M. acknowledges funding from the EMBL and the European Research Council starting grant (3DCellPhase^-^ 760067).

## Author contributions

X.Z. and J.M. conceived this work and wrote the manuscript. X.Z. performed data generation, processing and analysis, and prepared figures.

## References

1. Zhang, X., Sridharan, S., Zagoriy, I., Eugster Oegema, C., Ching, C., Pflaesterer, T., Fung, H.K.H., Becher, I., Poser, I., Muller, C.W., et al. (2023). Molecular mechanisms of stress-induced reactivation in mumps virus condensates. Cell 186, 1877–1894 e1827. 10.1016/j.cell.2023.03.015.

2. Zanetti, G., Prinz, S., Daum, S., Meister, A., Schekman, R., Bacia, K., and Briggs, J.A. (2013). The structure of the COPII transport-vesicle coat assembled on membranes. Elife 2, e00951. 10.7554/eLife.00951.

3. Hagen, W.J.H., Wan, W., and Briggs, J.A.G. (2017). Implementation of a cryo-electron tomography tilt-scheme optimized for high resolution subtomogram averaging. J. Struct. Biol. 197, 191–198. 10.1016/j.jsb.2016.06.007.

4. Mastronarde, D.N. (2005). Automated electron microscope tomography using robust prediction of specimen movements. J. Struct. Biol. 152, 36–51. 10.1016/j.jsb.2005.07.007.

5. Kremer, J.R., Mastronarde, D.N., and McIntosh, J.R. (1996). Computer visualization of three-dimensional image data using IMOD. J. Struct. Biol. 116, 71–76. 10.1006/jsbi.1996.0013.

6. Tegunov, D., and Cramer, P. (2019). Real-time cryo-electron microscopy data preprocessing with Warp. Nat. Methods 16, 1146–1152. 10.1038/s41592-019-0580-y.

7. Tegunov, D., Xue, L., Dienemann, C., Cramer, P., and Mahamid, J. (2021). Multi-particle cryo-EM refinement with M visualizes ribosome-antibiotic complex at 3.5 A in cells. Nat. Methods 18, 186–193. 10.1038/s41592-020-01054-7.

8. Castano-Diez, D., Kudryashev, M., Arheit, M., and Stahlberg, H. (2012). Dynamo: a flexible, user-friendly development tool for subtomogram averaging of cryo-EM data in high-performance computing environments. J. Struct. Biol. 178, 139–151. 10.1016/j.jsb.2011.12.017.

9. Nickell, S., Forster, F., Linaroudis, A., Net, W.D., Beck, F., Hegerl, R., Baumeister, W., and Plitzko, J.M. (2005). TOM software toolbox: acquisition and analysis for electron tomography. J. Struct. Biol. 149, 227–234. 10.1016/j.jsb.2004.10.006.

10. Forster, F., and Hegerl, R. (2007). Structure determination in situ by averaging of tomograms. Methods Cell Biol. 79, 741–767. 10.1016/S0091-679X(06)79029-X.

11. Zivanov, J., Nakane, T., Forsberg, B.O., Kimanius, D., Hagen, W.J., Lindahl, E., and Scheres, S.H. (2018). New tools for automated high-resolution cryo-EM structure determination in RELION-3. Elife 7. 10.7554/eLife.42166.

12. Pettersen, E.F., Goddard, T.D., Huang, C.C., Couch, G.S., Greenblatt, D.M., Meng, E.C., and Ferrin, T.E. (2004). UCSF Chimera--a visualization system for exploratory research and analysis. J. Comput. Chem. 25, 1605–1612. 10.1002/jcc.20084.

13. Pettersen, E.F., Goddard, T.D., Huang, C.C., Meng, E.C., Couch, G.S., Croll, T.I., Morris, J.H., and Ferrin, T.E. (2021). UCSF ChimeraX: Structure visualization for researchers, educators, and developers. Protein Sci. 30, 70–82. 10.1002/pro.3943.

14. Zheng, S., Wolff, G., Greenan, G., Chen, Z., Faas, F.G.A., Barcena, M., Koster, A.J., Cheng, Y., and Agard, D.A. (2022). AreTomo: An integrated software package for automated marker-free, motion-corrected cryo-electron tomographic alignment and reconstruction. J Struct Biol X 6, 100068. 10.1016/j.yjsbx.2022.100068.

15. Buchholz, T.O., Krull, A., Shahidi, R., Pigino, G., Jekely, G., and Jug, F. (2019). Content-aware image restoration for electron microscopy. Methods Cell Biol. 152, 277–289. 10.1016/bs.mcb.2019.05.001.

16. Rosenthal, P.B., and Henderson, R. (2003). Optimal determination of particle orientation, absolute hand, and contrast loss in single-particle electron cryomicroscopy. J. Mol. Biol. 333, 721–745. 10.1016/j.jmb.2003.07.013.

17. Hrabe, T., Chen, Y., Pfeffer, S., Cuellar, L.K., Mangold, A.V., and Forster, F. (2012). PyTom: a python-based toolbox for localization of macromolecules in cryo-electron tomograms and subtomogram analysis. J. Struct. Biol. 178, 177–188. 10.1016/j.jsb.2011.12.003.

18. Chaillet, M.L., van der Schot, G., Gubins, I., Roet, S., Veltkamp, R.C., and Forster, F. (2023). Extensive Angular Sampling Enables the Sensitive Localization of Macromolecules in Electron Tomograms. Int. J. Mol. Sci. 24. 10.3390/ijms241713375.

19. Chen, M., Bell, J.M., Shi, X., Sun, S.Y., Wang, Z., and Ludtke, S.J. (2019). A complete data processing workflow for cryo-ET and subtomogram averaging. Nat. Methods 16, 1161–1168. 10.1038/s41592-019-0591-8.

20. Poge, M., Mahamid, J., Imanishi, S.S., Plitzko, J.M., Palczewski, K., and Baumeister, W. (2021). Determinants shaping the nanoscale architecture of the mouse rod outer segment. Elife 10. 10.7554/eLife.72817.

21. Roy, A., Kucukural, A., and Zhang, Y. (2010). I-TASSER: a unified platform for automated protein structure and function prediction. Nat. Protoc. 5, 725–738. 10.1038/nprot.2010.5.

22. Jumper, J., Evans, R., Pritzel, A., Green, T., Figurnov, M., Ronneberger, O., Tunyasuvunakool, K., Bates, R., Zidek, A., Potapenko, A., et al. (2021). Highly accurate protein structure prediction with AlphaFold. Nature 596, 583–589. 10.1038/s41586-021-03819-2.

23. Liebschner, D., Afonine, P.V., Baker, M.L., Bunkoczi, G., Chen, V.B., Croll, T.I., Hintze, B., Hung, L.W., Jain, S., McCoy, A.J., et al. (2019). Macromolecular structure determination using X-rays, neutrons and electrons: recent developments in Phenix. Acta Crystallogr. D Struct. Biol. 75, 861–877. 10.1107/S2059798319011471.

24. Emsley, P., Lohkamp, B., Scott, W.G., and Cowtan, K. (2010). Features and development of Coot. Acta Crystallogr. D Biol. Crystallogr. 66, 486–501. 10.1107/S0907444910007493.

25. de Teresa-Trueba, I., Goetz, S.K., Mattausch, A., Stojanovska, F., Zimmerli, C.E., Toro-Nahuelpan, M., Cheng, D.W.C., Tollervey, F., Pape, C., Beck, M., et al. (2023). Convolutional networks for supervised mining of molecular patterns within cellular context. Nat. Methods 20, 284–294. 10.1038/s41592-022-01746-2.

26. Watanabe, R., Buschauer, R., Bohning, J., Audagnotto, M., Lasker, K., Lu, T.W., Boassa, D., Taylor, S., and Villa, E. (2020). The In Situ Structure of Parkinson’s Disease-Linked LRRK2. Cell 182, 1508–1518 e1516. 10.1016/j.cell.2020.08.004.

27. Zivanov, J., Oton, J., Ke, Z., von Kugelgen, A., Pyle, E., Qu, K., Morado, D., Castano-Diez, D., Zanetti, G., Bharat, T.A.M., et al. (2022). A Bayesian approach to single-particle electron cryo-tomography in RELION-4.0. Elife 11. 10.7554/eLife.83724.

28. Wan, W., Kolesnikova, L., Clarke, M., Koehler, A., Noda, T., Becker, S., and Briggs, J.A.G. (2017). Structure and assembly of the Ebola virus nucleocapsid. Nature 551, 394–397. 10.1038/nature24490.

29. Wang, Z., Grange, M., Pospich, S., Wagner, T., Kho, A.L., Gautel, M., and Raunser, S. (2022). Structures from intact myofibrils reveal mechanism of thin filament regulation through nebulin. Science 375, eabn1934. 10.1126/science.abn1934.

